# Molecular architecture of colossal surface layers from hyperthermophilic archaea

**DOI:** 10.64898/2026.08.21.746120

**Authors:** Ido Caspy, Virginija Cvirkaite-Krupovic, Sofie van Dorst, Andriko von Kügelgen, Zephyr Ford, Vikram Alva, Mart Krupovic, Tanmay A. M. Bharat

## Abstract

Surface layers (S-layers) are paracrystalline protein lattices that form the outermost layer of the cell envelope in most archaea, providing structural support, protecting against external insults, and co-ordinating interactions with their environment. Despite their widespread occurrence, the molecular and structural details of S-layer architecture in hyperthermophilic archaea remain largely unknown. Here, we report the structure and cellular architecture of the S-layer from the hyperthermophilic archaeon *Pyrobaculum arsenaticum* by combining *in situ* electron cryotomography with single-particle electron cryomicroscopy, AlphaFold modelling, and peptide-fingerprinting mass spectrometry. We show that the S-layer is formed by an uncharacterised 292-kDa S-layer protein (SLP) extending 37 nm from the cytoplasmic membrane, making it, to our knowledge, the largest SLP structurally characterised to date. This SLP has a remarkable multidomain architecture comprising 19 immunoglobulin-like domains, 14 canonical and five non-canonical, organised into a lattice-forming core, a stalk, and a unique crown domain that stabilise the S-layer. Comparative genomic analyses unearthed homologous colossal SLP candidates across Thermoproteota, indicating that this distinctive architecture is conserved across diverse archaeal lineages. Together, our findings provide a structural framework for understanding the cell-surface organisation in hyperthermophilic archaea and suggest that these colossal S-layers represent a specialised adaptation to life at high temperatures.

## Introduction

Surface layers (S-layers) are two-dimensional, paracrystalline arrays made of either protein or glycosylated protein subunits, called surface layer proteins (SLPs), which form the outermost cell-envelope component in many bacteria and archaea^1–3^. These highly ordered assemblies are formed through the self-assembly of SLPs into curved sheets with different planar symmetries^4,5^. Since most archaea lack the typical peptidoglycan cell wall found in monoderm (also known as Gram-positive) bacteria or the second membrane found in diderm (also known as Gram-negative) bacteria, S-layers are the major structural component scaffolding the cytoplasmic membrane. Therefore, archaeal S-layers play fundamental roles in maintaining cellular integrity, architecture, and morphology^2,6,7^. Beyond their structural function, S-layers contribute to a wide range of biological processes, including the molecular sieving of metabolically relevant, scarce molecules^8^, protection from environmental insults such as high temperature^9^ or osmotic pressure^10,11^, and the mediation of cell-cell interactions^12,13^.

SLPs typically possess complex, multidomain architectures that support varied structural and functional roles within a single polypeptide^4^. Apart from the two major structural requirements of lattice formation and cellular anchoring, specific SLP domains may be glycosylated, a modification that is likely to contribute to the S-layer stability and play a key role in intercellular communication, allowing species-specific self-versus-non-self discrimination^3,14,15^. The evolution of such multidomain proteins is widely thought to occur through mechanisms such as gene duplication, domain fusion, and subsequent functional diversification, processes that facilitate the emergence of novel protein functions^16–18^. Such evolutionary mechanisms may have played a significant role in shaping the diversity and adaptability of bacterial and archaeal SLPs.

Many extremophilic archaea prosper in habitats that are inhospitable to most other forms of life. These extremophiles thrive under conditions of high temperature, extreme pH, high salinity, elevated pressure, and even combinations of these extreme physical conditions, requiring specialised molecular adaptations to sustain the most basic cellular functions^19,20^. Thermophilic and hyperthermophilic archaea, in particular, have evolved mechanisms that allow their cells and cell envelope to remain functional at temperatures exceeding those tolerated by mesophilic organisms^21^. Protein thermostability can be achieved by employing a variety of structural adaptations, including increased numbers of disulfide bonds, enhanced salt-bridge networks, hydrophobic-core packing, glycosylation, and increased surface-charge interactions^22,23^. The combined effect of these individual adaptations stabilises protein structure under thermal stress while maintaining biological activity under conditions that would otherwise compromise macromolecular integrity or functionality.

Given their exposed cellular location and essential structural role, SLPs from thermophilic archaea provide an interesting system in which to investigate how S-layers are arranged in organisms living in extreme environments. However, despite extensive biochemical and genomic characterisation of archaeal S-layers, information from hyperthermophilic organisms remains limited^24–28^. In particular, high-resolution structural studies of SLPs from thermophilic organisms have historically been scarce^29^, and no S-layer structures from hyperthermophilic prokaryotes, defined as organisms with an optimal growth temperature above 80 °C, are available, leaving significant gaps in our understanding of how these proteins maintain stability and assembly under severe thermal stress. Elucidating the structural features that underpin thermostability in archaeal SLPs is therefore important for understanding both the evolution of extremophilic adaptations and the fundamental principles governing cell envelope organisation in hyperthermophilic archaea.

To address these knowledge gaps, we studied the hyperthermophilic crenarchaeon *Pyrobaculum arsenaticum*, which was isolated from a hot spring and grows optimally at temperatures approaching 95 °C^30,31^. Together with other members of the *Pyrobaculum* genus, *P. arsenaticum* has been shown to harbour an S-layer that serves as its primary cell envelope component^32–36^. Here, we examined the S-layer architecture of *P. arsenaticum* and the related hyperthermophile *Pyrobaculum oguniense* by electron cryotomography (cryo-ET), revealing a conserved cell envelope organisation *in situ*. We then used subtomogram averaging (STA) of the *P. arsenaticum* S-layer to define its native architecture. Next, we solved the S-layer structure at 5.7 Å-resolution using electron cryomicroscopy single-particle analysis (cryo-EM SPA), which, combined with AlphaFold^37,38^ modelling and mass spectrometry allowed us to identify its colossal (292 kDa) SLP. Finally, comparative genomic analyses and structure predictions identified homologous colossal SLPs across three classes of the phylum Thermoproteota—Thermoprotei (Thermoproteales and Thermofilales), Nitrososphaeria_A (Caldarchaeales), and Methanomethylicia (Culexarchaeales)— indicating that this distinctive architecture is conserved across multiple hyperthermophilic lineages. Collectively, our data show how a colossal SLP, the largest SLP structurally characterised to date, forms the *Pyrobaculum* S-layer, revealing clues to its assembly and thermostability, while also illuminating molecular details of cell envelope organisation in hyperthermophilic archaea.

## Results

### Cellular architecture of the S-layer

We first performed cryo-ET on two species from the *Pyrobaculum* genus, *P. arsenaticum* and *P. oguniense*, to shed light on S-layer composition and architecture *in situ* (Methods). Analysis of the cellular cryo-ET data showed that both *P. arsenaticum* and *P. oguniense* cells were coated with an S-layer extending nearly ∼37 nm from the cytoplasmic membrane (Fig. 1A-B, Supplementary Movies 1 and 2). Consistent with previous reports^32^, the S-layer appeared to be organised into a hexagonal lattice, with a centre-to-centre spacing of ∼30 nm between neighbouring hexamers (Fig. 1C-D). Several structural features of the S-layer were observed directly in the reconstructed tomograms. These included elongated, pillar-like densities extending from the cell membrane to the lattice-forming region of the S-layer and a large pore-like density formed at the centre of the S-layer hexamer along the pillar-like densities (Fig. 1C-F). Intriguingly, we identified gaps in the S-layer lattice in both *P. arsenaticum* and *P. oguniense* (Fig. 1D and Supplementary Fig. 1). In previously characterised archaeal hexagonal S-layers, some S-layers undergo an oligomeric transition near the cell poles, forming pentameric or heptameric defects, to accommodate the increased surface curvature, while ensuring continuous coverage by the S-layer lattice^8,39^. In *Pyrobaculum*, by contrast, the lattice is discontinuous, with gaps observed at the cell poles to accommodate the changing curvature in these regions. Gaps in S-layer lattices have previously been observed in the diderm bacteria *Caulobacter crescentus*^40^ and *Corynebacterium glutamicum*^41,42^.

**Fig. 1.**
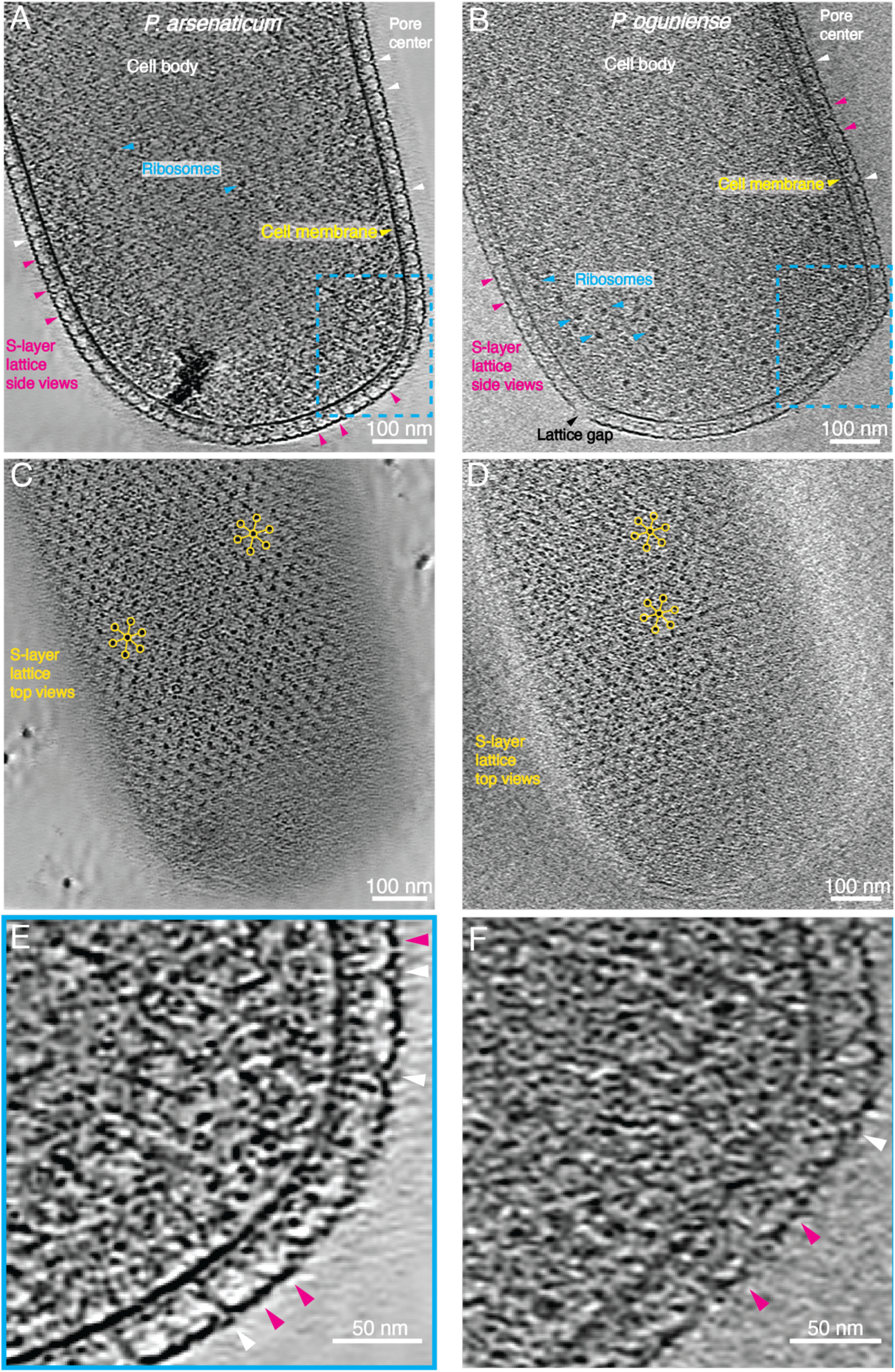
- Cryo-ET of two *Pyrobaculum* species. Tomographic slices showing the cell body of *P. arsenaticum* (**A**) and *P. oguniense* (**B**). The cell membrane (yellow arrowheads), ribosomes (cyan arrowheads), S-layer side views (magenta arrowheads), a pore-like density at the hexamer centre (white arrowheads) and a gap in the S-layer lattice (black arrowhead) are marked. *P. arsenaticum* (**C**) and *P. oguniense* (**D**) tomograhic slices showing a top view of the hexagonal S-layer lattice coat on the cell surface. Hexameric S-layer subunits are marked with yellow circles. Zoomed-in views of the *P. arsenaticum* (**E**) and *P. oguniense* (**F**) cell edge highlighting the S-layer lattice side views (magenta arrowheads) and pore-like density (white arrowheads).

To decipher the organisation of the S-layer, we performed STA directly on the cell surface of *P. arsenaticum* from whole-cell cryo-ET (shown in Fig. 1). The resulting 18 Å-resolution STA map highlighted prominent features of the S-layer that were observed in raw tomograms above, such as the pillar-like and crown-like densities (Fig. 2A). To resolve the lattice arrangement, we performed STA refinements with larger box sizes to include the six proximal hexamers surrounding the central hexamer (Fig. 2B) and then mapped their locations back onto the tomograms. The resulting mapping of the S-layer subtomograms confirmed that it is arranged as an imperfect hexagonal lattice on the cell surface, with slight variations in the spacing between the hexamers in the lattice plane (Fig. 2C-F). These lattice imperfections are consistent with the observed gaps in the lattice (Supplementary Fig. 1), which were confirmed by the STA analysis.

**Fig. 2.**
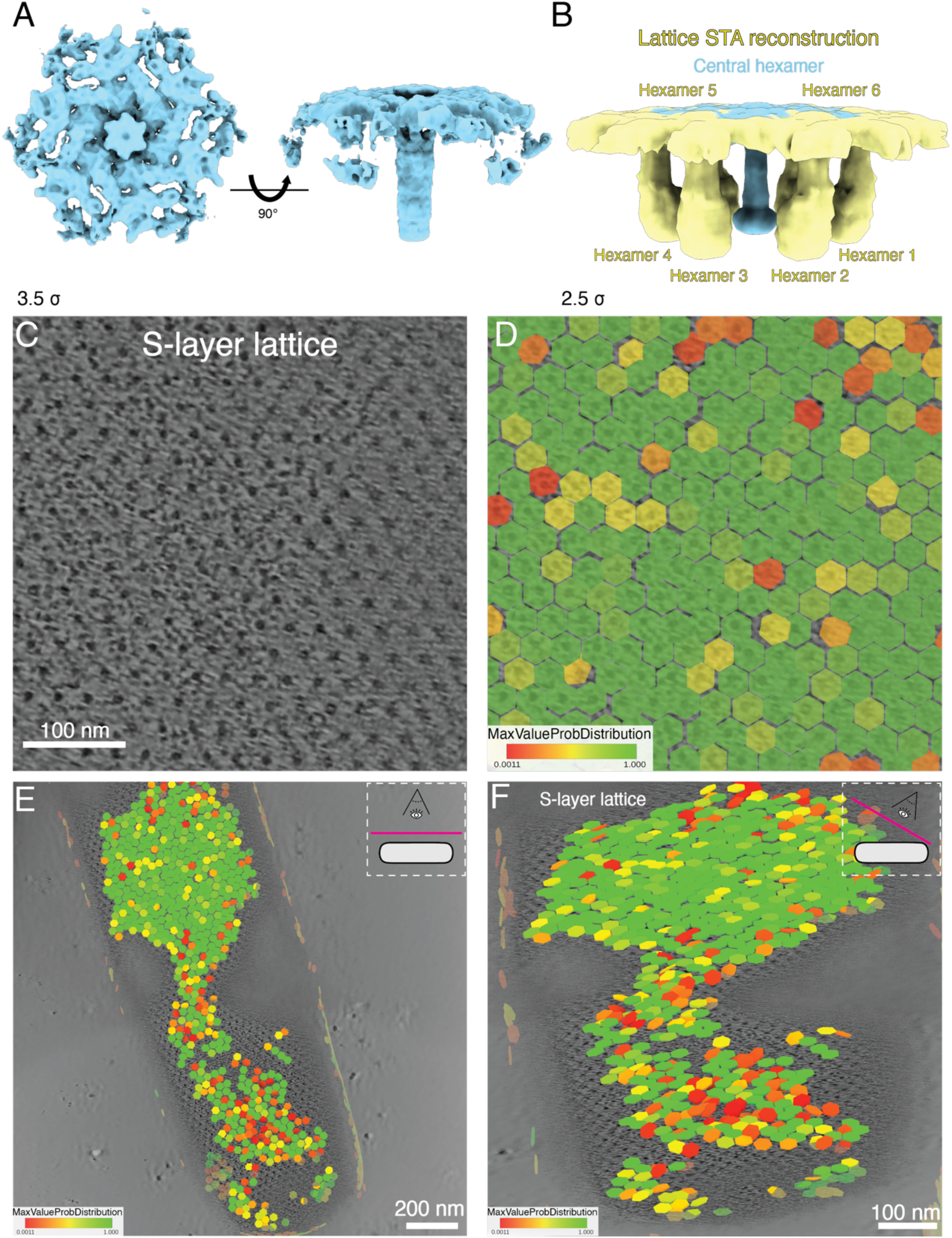
- *P. arsenaticum* cryo-ET STA reveals the *in situ* S-layer lattice arrangement. **A**) Subtomogram averaging (STA) reconstruction of the *P. arsenaticum* S-layer hexamer shown in top and side views. **B**) STA with a larger box around the central hexamer, surrounded by six bordering hexamers. **C**) Zoom-in of a tomographic slice displaying the S-layer arrangement from a top view. The dot-shaped densities are orthogonal views of the pillar-like densities seen the STA map shown in panels A-B. **D**) Plotting the STA final particle positions after alignment onto the same tomographic slice as in panel C. Particles are presented as transparent hexagons and coloured according to the RELION5^89^ variable *MaxValueProbDistribution* indicating per-particle orientation confidence score. **E-F**) Zoomed-out views of the lattice map showing all S-layer particles identified in the tomogram overlaid onto the tomographic slice, presented in the same colour scheme as (D). The inset describes the tomogram oblique slice viewing angle with respect to the cell orientation.

### Cryo-EM structure of the S-layer

Next, to understand the molecular arrangement of the SLP within the S-layer, we purified cell envelopes of *P. arsenaticum* cells and applied cryo-EM SPA to this specimen, following previous successful application of this technique to S-layers^39,41^. Using this approach, we resolved a 5.75 Å-resolution cryo-EM map (Fig. 3, Supplementary Figs. 2-3, and Supplementary Table 1), which could not be improved by increasing the data set size or by extensive classification of different states in image processing. Our data suggest an innate flexibility in both the lattice-forming region and the pillar, limiting the resolution of the final reconstruction. While at this resolution we could not discern the individual side-chain densities, upon closer inspection many secondary structure elements were identifiable in the cryo-EM map, which we subsequently utilised for fitting an atomic model.

**Fig. 3.**
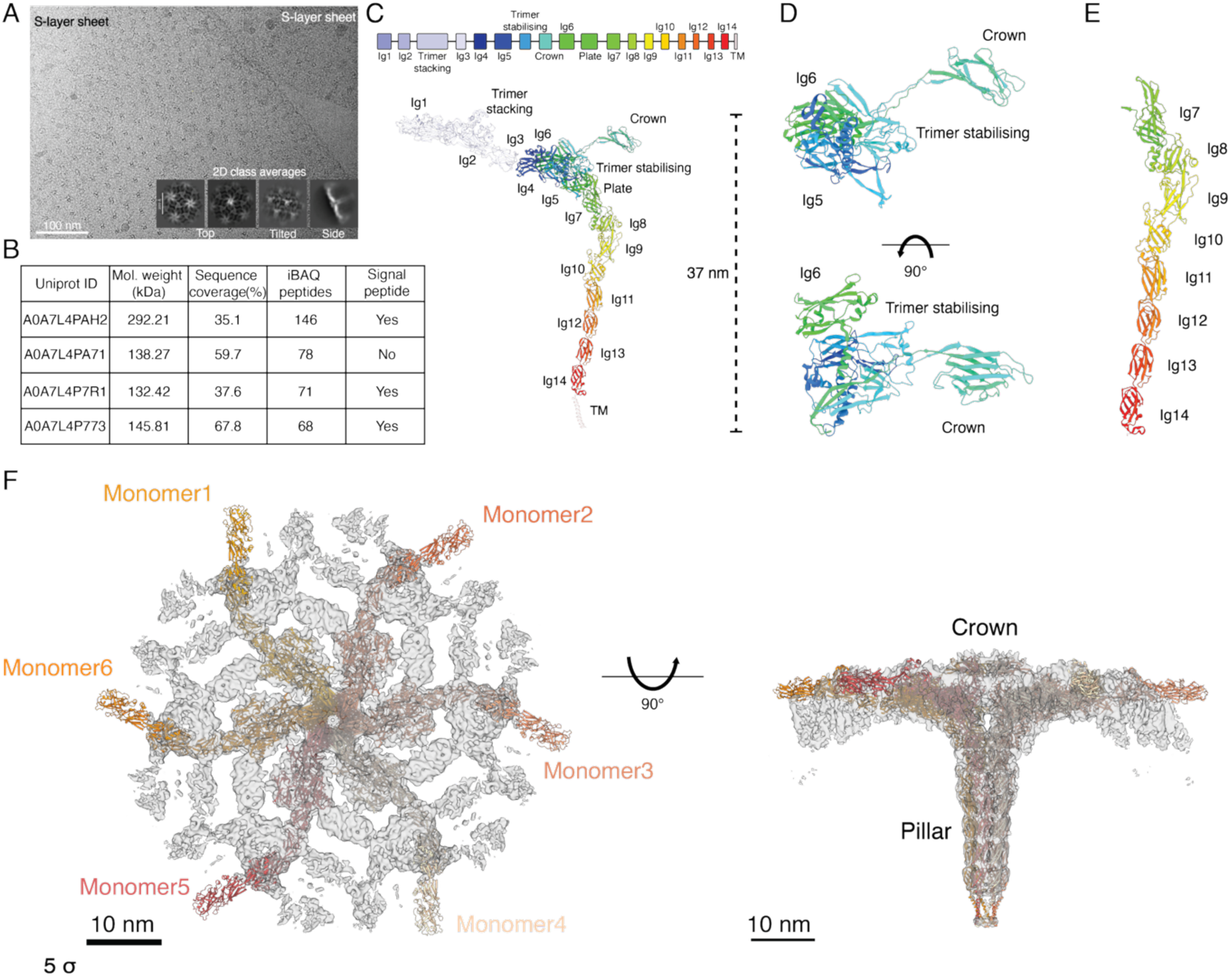
- Cryo-EM SPA structure determination, protein identification and model building of PySLP. **A**) Representative cryo-electron micrograph displaying two purified PySLP S-layer sheets (marked). The insets show two-dimensional class averages of the S-layer in various orientations. **B**) Top four hits from peptide fingerprinting mass-spectrometry of purified cell envelopes of *P. arsenaticum* (iBAQ peptides - intensity-based absolute quantification, calculated by summing the intensity peaks of all detected peptides and dividing by the theoretical peptide number of a specific protein). **C**) Domain arrangement of the PySLP monomer, the size of each box is to scale with the number of residues in each domain. The light tones of domains Ig1, Ig2, Ig3, trimer stacking and the TM helix represent lower resolution regions in our cryo-EM map. **D**) Zoom-in on Ig5, trimer stabilising, crown, and Ig6 domains, viewed from two orthogonal orientations. **E**) Zoom-in on the C-terminal tandem domains Ig7-Ig14. **F**) Hexameric PySLP model fitted into the cryo-EM 5.75-Å resolution map (transparent dark grey) viewed from top (left) and membrane plane (right).

As the identity and sequence of the SLP were unknown, we used peptide-fingerprinting mass-spectrometry (MS) of the cell envelope preparations to identify possible candidates. In the analysis of the MS data, we sought to identify candidate proteins that satisfied three criteria: (i) a large molecular weight, consistent with the unusually bulky S-layer observed in the density map, (ii) high abundance within the sample, as SLPs are typically among the most highly expressed proteins in cells that harbour them^5^, and (iii) the presence of an N-terminal signal peptide to enable secretion across the cytoplasmic membrane. Based on these criteria, we identified four initial candidate proteins that met at least two of these requirements (Fig. 3B). Mass spectrometry analysis identified a putative SLP encoded by the gene *HC235_06945* (UniProtKB A0A7L4PAH2). This uncharacterised protein was both the most abundant hit in the sample and the protein with the largest predicted molecular weight, exceeding 292 kDa (2,667 residues; Fig. 3B). Notably, HC235_06945 is also the largest protein encoded in the *P. arsenaticum* genome. Moreover, the first 23 amino acids were predicted to constitute a signal peptide^43^, providing additional support for its assignment as the SLP.

As a first step, we attempted to predict the entire protein using AlphaFold3^44^. However, the output generated by AlphaFold3 (Supplementary Fig. 4) could not be fitted satisfactorily into the cryo-EM map. We therefore divided the protein sequence into smaller, roughly equal-sized segments that could be docked into the cryo-EM map (Fig. 3C, Supplementary Fig. 5). The structures of the N-terminal segment, encompassing five domains, including a tightly associated domain pair (residues 24–828), the middle segment, comprising six domains (residues 829–1835), and the C-terminal segment, comprising the final eight domains (residues 1836–2659), were predicted using the ColabFold^38^ implementation of AlphaFold2^37^ (Fig. 3D-E, Supplementary Fig. 5). The predicted segments fit well within our experimental cryo-EM map and could be used for restrained model-to-map refinement (Supplementary Table 1). Many secondary-structure elements resolved in the map aligned well with the refined model (Supplementary Fig. 6). The only exception was the crown domain (residues 1312–1461), for which we generated a separate prediction using six copies of the crown polypeptide (Supplementary Fig. 5), consistent with our observation that the domain is located at the centre of the hexamer. After refining each part of the polypeptide separately, we merged the models to construct a complete model of a single SLP, which was subsequently copied to build the entire S-layer hexamer (Fig. 3F, Supplementary Movie 3).

Using this hexameric model, we then re-examined the cryo-ET STA map (from Fig. 2) to verify that the S-layer arrangement observed *in situ* matched our cryo-EM SPA map and the atomic model of the hexamer. A comparison of the maps and the model at the resolution of the STA map (18 Å) showed a close match, demonstrating that the native S-layer had been correctly reconstructed (Fig. 4). This comparison also illustrated that the architecture observed on the native cell surface was maintained in the purified S-layer. As a further check, we investigated whether another abundant candidate identified by peptide-fingerprinting MS could account for our cryo-EM map. We used AlphaFold2 to obtain a model of this second largest protein (UniProtKB A0A7L4P773) (Fig. 3B, Supplementary Fig. 7). Fitting the model into the map revealed that large portions of the cryo-EM density remained unexplained, and the addition of further copies could not account for all of the density in the map. In particular, the densities corresponding to the pillar, crown, and the middle lattice-forming plate domain could not be explained by this protein or by the other abundant candidates in our list. These results strongly support the assignment of A0A7L4PAH2, since it is the only protein that can explain the cryo-EM density of the lattice; hereafter, we refer to it as PySLP (*Pyrobaculum* SLP).

**Fig. 4.**
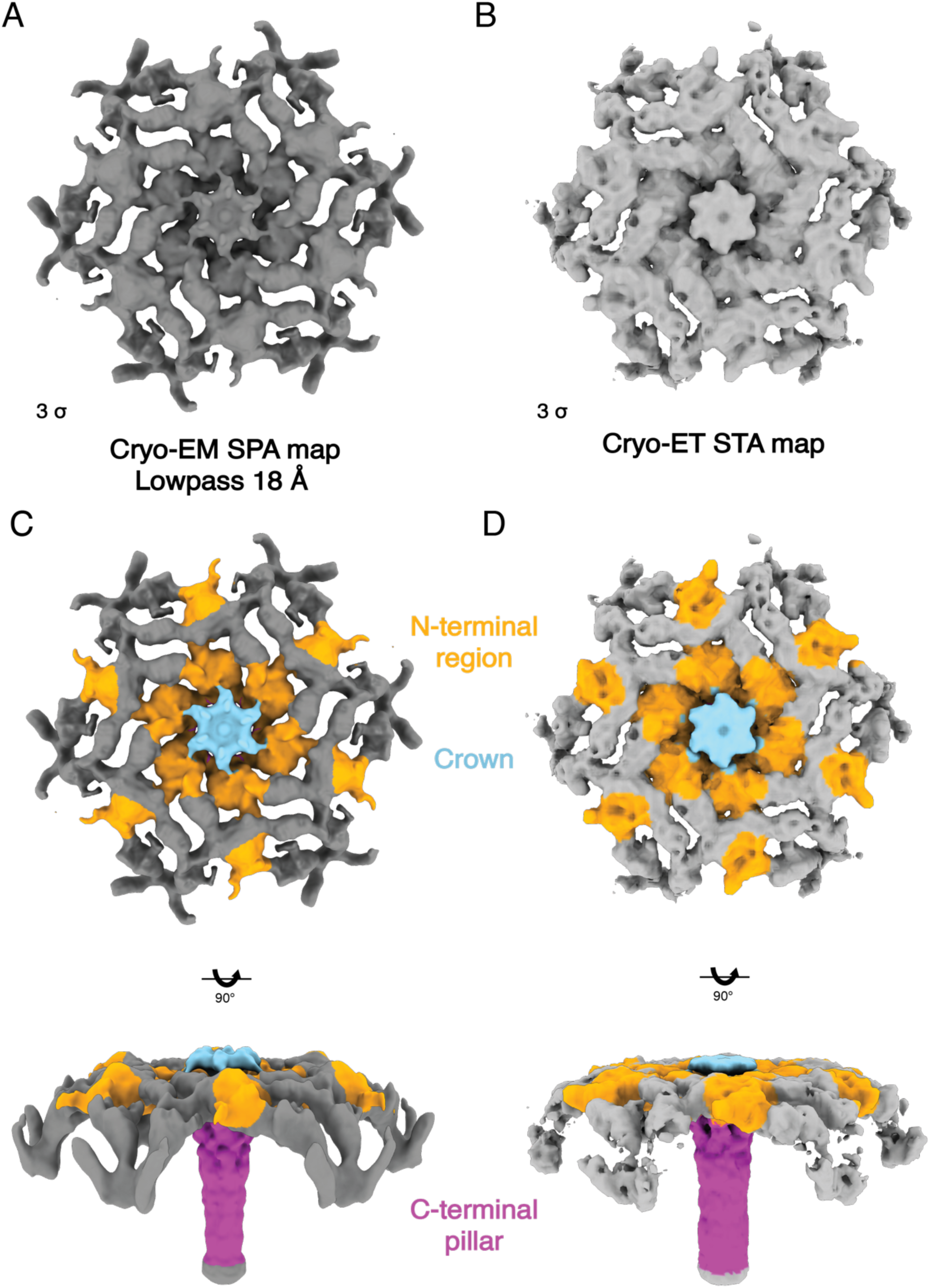
- Comparison of the cryo-EM SPA and cryo-ET STA map. Cryo-EM SPA (**A**) and cryo-ET STA (**B**) maps lowpass filtered to 18 Å. Same maps as in panels A-B, but coloured according to PySLP domains resolved in the SPA (**C**) and STA (**D**) maps. The N-terminal region is coloured in orange, crown in cyan, and C-terminal pillar in purple. The colourised maps are shown in two orthogonal orientations.

As previously observed in other S-layers^8,45^, immunoglobulin (Ig)-like folds constitute the predominant tertiary structural motif identified in PySLP. Overall, PySLP contains 19 Ig-like domains, of which 14 adopt canonical Ig-like folds (Ig1–Ig14), whereas the remaining five are more divergent and/or contain extensive insertions. For example, the third N-terminal domain, referred to as the trimer-stacking domain, comprises two tightly packed Ig-like folds with extensive insertions. Owing to their intimate association, we treated these folds as a single architectural domain. The middle region displays further topological complexity, with Ig6, the plate domain, and the crown domain all inserted within the trimer-stabilising domain. Together, the 19 domains form the lattice-forming region, including Ig1-Ig6 and the additional non-canonical domains, and the pillar (Ig7-Ig14), extending from the lattice layer through to the cell membrane (Fig. 3F). At the C-terminal region, both AlphaFold2 and DeepTMHMM^46^ predicted the presence of a transmembrane helix, which we propose anchors the S-layer to the cytoplasmic membrane, as previously suggested for *Pyrobaculum aerophilum* based on the analysis of ultrathin sections^47^ and described for other archaeal SLPs^7,8,39,48^.

We next examined the cryo-EM map for evidence of potential glycosylation. Although the resolution of the cryo-EM map was insufficient to resolve individual side chains or glycans, we identified several prominent densities (Supplementary Fig. 8A) located near asparagine (Asn) residues found within the consensus N-glycosylation motif (Asn-X-Ser/Thr). The shape and localisation of these densities were consistent with glycan moieties previously described in *Pyrobaculum* pilus structures^49,50^, although confident assignment of these densities as glycans was not possible at this resolution. Inspection of the charge distribution on PySLP revealed that unlike previously resolved SLPs^8,51,52^, the surface exposed to the extracellular environment, especially the central area of the hexamer composed of the six crown domains, was strongly positively charged (Supplementary Fig. 8B). We speculate that these electropositive regions may contribute to the binding of negatively charged ions, such as arsenate and selenate, which serve as terminal electron acceptors in *P. arsenaticum* respiration^30^.

The pronounced divergence of the non-canonical Ig-like domains was also reflected in the limited number of homologous sequences identified in the AlphaFold2-based multiple sequence alignments (Supplementary Fig. 9). The N-terminal region spanning residues 340-828 and the crown domain (residues 1312-1461) each produced only a handful (20-30) sequence matches, in contrast to hundreds of homologous sequences identified for the canonical Ig-like domains of the C-terminal pillar (Supplementary Fig. 9C). Structure-based searches using DALI^53^ and FoldSeek^54^, together with sequence-based searches using HHpred^55^, did not produce significant matches for the crown domain, suggesting that it represents a highly divergent Ig-like fold. At the hexagonal interface, six copies of the crown domain assemble into a crown-like structure elevated 5 nm above the lattice-forming region. Separately, six segments, each comprising the six C-terminal domains of one PySLP, associate to form a central channel within the pillar that constricts to 12 Å and is lined with charged residues. This channel could permit the passage of solutes and small molecules, and potentially single-stranded DNA.

This colossal PySLP represents the largest SLP structure experimentally described to date, extending 37 nm from the cell surface, i.e. from the putative membrane-associated transmembrane helix to the lattice-forming region (Fig. 3C). To elucidate the organisation of the fully assembled S-layer, we performed rigid-body fits of the hexameric model into the cryo-EM map and constructed a larger section of the lattice by placing six neighbouring hexamers around the central hexamer (Fig. 5). Although the resolution of the map is not sufficient to resolve interaction details at the side chain level, the resulting lattice model accounts for all the densities in our cryo-EM map, and shows how this exceptionally large S-layer is stabilised chiefly by trimeric interfaces formed between three adjacent hexamers, where the stacking domains sit one on top of another. These interactions are further supported by contacts between the N-terminal Ig1 domains and the neighbouring trimer-stacking domain (Fig. 3C, 5, and 6A, Supplementary Movie 3). Comparing the lattice model with a tomographic slice from a *P. arsenaticum* cell confirmed a cytoplasmic membrane-to-lattice distance of approximately 37 nm in the native cellular context (Fig. 6B).

**Fig. 5.**
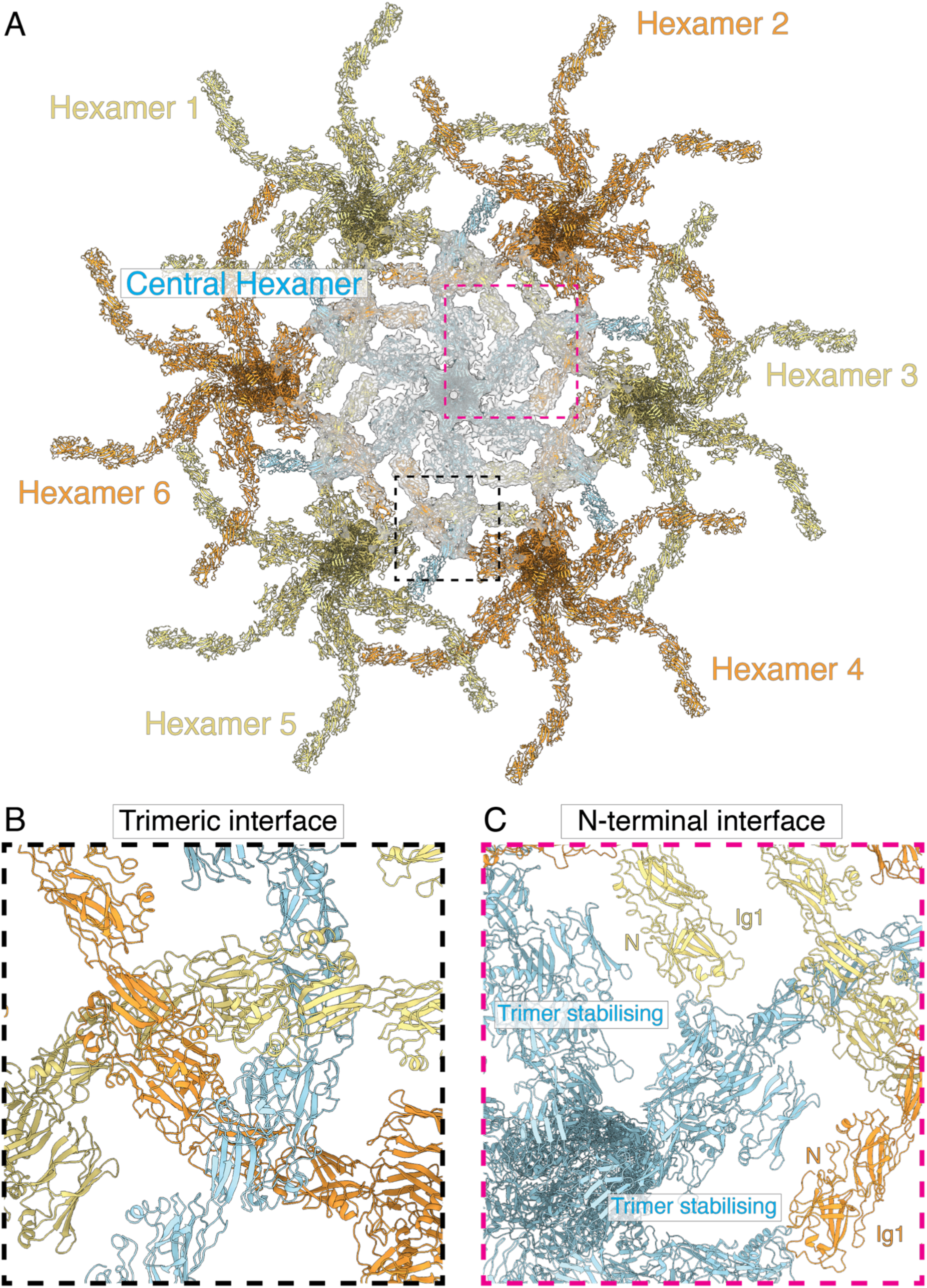
- Lattice model fit to the cryo-EM SPA map and predicted inter hexamer interfaces. **A)** Lattice model in ribbon representation containing the central hexamer with its six adjacent hexamers rigid body overlaid onto the cryo-EM map (clash score 14.45). **B**) Trimeric stacking interface generated by three neighbouring hexamers. **C**) Predicted trimer stabilising domain interface with domain Ig1 in the N-terminus of an adjacent hexamer.

**Fig. 6.**
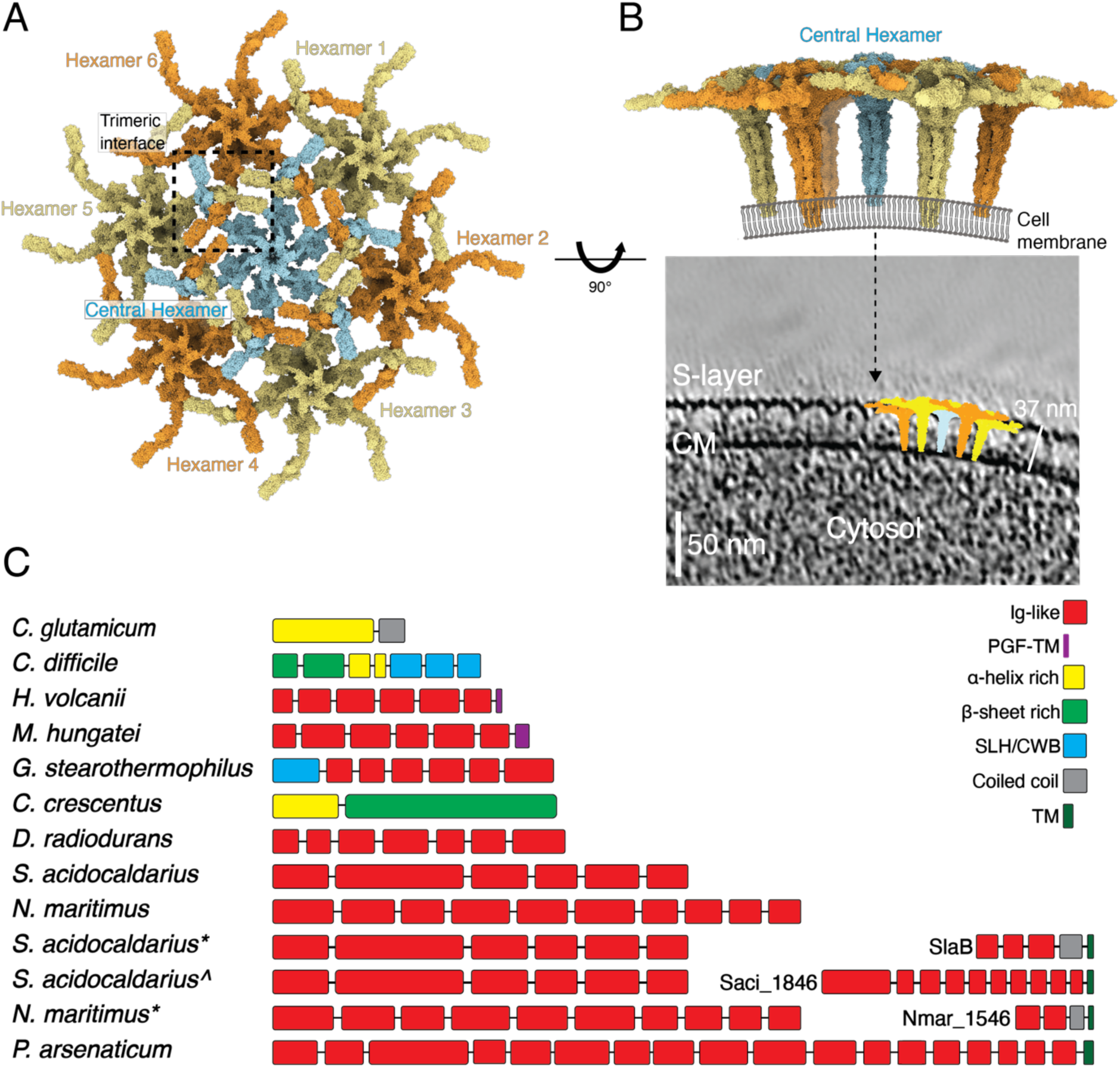
- PySLP lattice model comparison with cryo-ET and domain comparison with other SLPs. **A)** Complete lattice arrangement PySLP (top view is shown, observed from outside the cell). **B)** Top - Side view of the lattice arrangement with a cartoon of cell membrane added. Bottom - The lattice model containing five hexamers overlaid onto a tomographic slice. The 37 nm distance from the membrane to the edge of the S-layer is marked. **C**) Comparative analysis of the overall SLP size and corresponding domain arrangement of representative, experimentally characterised divergent SLP structures. Monoderm bacteria - *Clostridium difficile*^57^ (PDB ID 7ACY), *Geobacillus stearothemophilus*^52^ (4AQ1); diderm bacteria - *Caulobacter crescentus*^40^ (6Z7P); *Deinococcus radiodurans*^45^ (8CKA), *Corynebacterium glutamicum*^41^ (9HPN), and archaea - *Haloferax volcanii*^39^ (7PTR), *Methanospirillum hungatei*^56^, *Sulfolobus acidocaldarius*^29^ (7ZCX) and *Nitrosopumilus maritimus*^8^ (8C8L). *S. acidocaldarius* is shown together with two of its cell-anchoring proteins, SlaB* (8QOX)^29^ and Saci_1846^ (UniProtKB Q4J7S9)^7^, and *N. maritimus*\* with its SlaB homologue Nmar_1546 (UniProtKB A9A4Y8)^8^. The domain length is scaled with the number of residues, and colour coded with respect to its tertiary structure.

Comparison of the *P. arsenaticum* SLP with other experimentally determined high-resolution S-layer structures (Fig. 6C) demonstrates that Ig-like domains constitute a recurrent structural motif across a broad range of archaeal and bacterial SLPs^8,29,39–42,45,52,56,57^, although they are not universally present. The abundance of tandem Ig-like domains found in SLPs across diverse prokaryotic lineages has been suggested to arisen through repeated domain duplication and subsequent divergence during evolution^58^. Examining the domain architecture of experimentally characterised Ig-rich archaeal SLPs further highlights the exceptional size and complexity of PySLP. Whereas the SLPs of *Haloferax volcanii* and *Nitrosopumilus maritimus* contain six and ten Ig-like domains, respectively, PySLP contains 19 Ig-like domains (14 canonical and 5 non-canonical). This marked variation suggests that expansion and contraction of Ig-like domain arrays have played an important role in the diversification of archaeal S-layers. Collectively, these comparisons identify PySLP as the largest and most domain-rich SLP structurally described to date (Fig. 6C).

### Widespread occurrence of PySLP-like S-layers across Thermoproteota

Homologous, uncharacterised proteins of similar length were identified by BLAST^59^ across all five genera within the order *Thermoproteales*, most prominently among *Pyrobaculum* species, but also in *Thermoproteus*, *Vulcanisaeta*, *Thermocaldium*, and *Caldivirga* species^60^. Previous studies of *Thermoproteus* S-layers suggested a similar lattice morphology and arrangement, although the identity of the SLP remained undetermined^24,25^. The *T. tenax* gene *TTX_1887* had previously been proposed to encode the SLP^61^. We segmented representative candidate SLP sequences, including TTX_1887 in the same manner as the PySLP sequence and predicted their structures using AlphaFold3^44^. Predicted structures were generated for candidate SLPs from *Thermoproteus tenax* (UniProtKB G4RLQ9), *Thermocladium modestius* (UniProtKB A0A830GSK0), *Vulcanisaeta distributa* (UniProtKB E1QP79), and *Caldivirga maquilingensis* (UniProtKB A8M9H4). Comparison of these predicted models with our *P. arsenaticum* structure revealed broadly similar architectures, except for the absence of the crown domain (Fig. 7A). Indeed, the crown, with its positively charged surface, appear to be a distinguishing feature of the *Pyrobaculum* SLPs, suggesting functional specialisation relative to SLPs of other *Thermoproteales*. AlphaFold models further predicted a conserved β-hairpin between the final Ig-like domain and the transmembrane helix in all examined homologs. Although the cryo-EM density in this part of the map is of lower resolution, in the fitted PySLP hexamer, the β-hairpin from each SLP projects towards the central axis and positions a phenylalanine side chain into the channel lumen, forming a putative ring of six aromatic residues that may constitute a hydrophobic constriction. Using the above-mentioned representative sequences as queries, we identified additional homologous proteins in annotated *Thermoproteales* species from each genus and generated a multiple sequence alignment. The alignment revealed conserved regions, particularly within the Ig-like domains and the transmembrane helix. The extreme C-terminal cytoplasmic tail immediately following the transmembrane helix also showed conservation, being highly basic in all examined homologs. Notably, all examined homologs terminate in lysine, while the preceding tail residues display lineage-specific conservation. Together, these sequence and structural features support a common evolutionary origin and conserved membrane-anchoring architecture for this protein family.

**Fig. 7.**
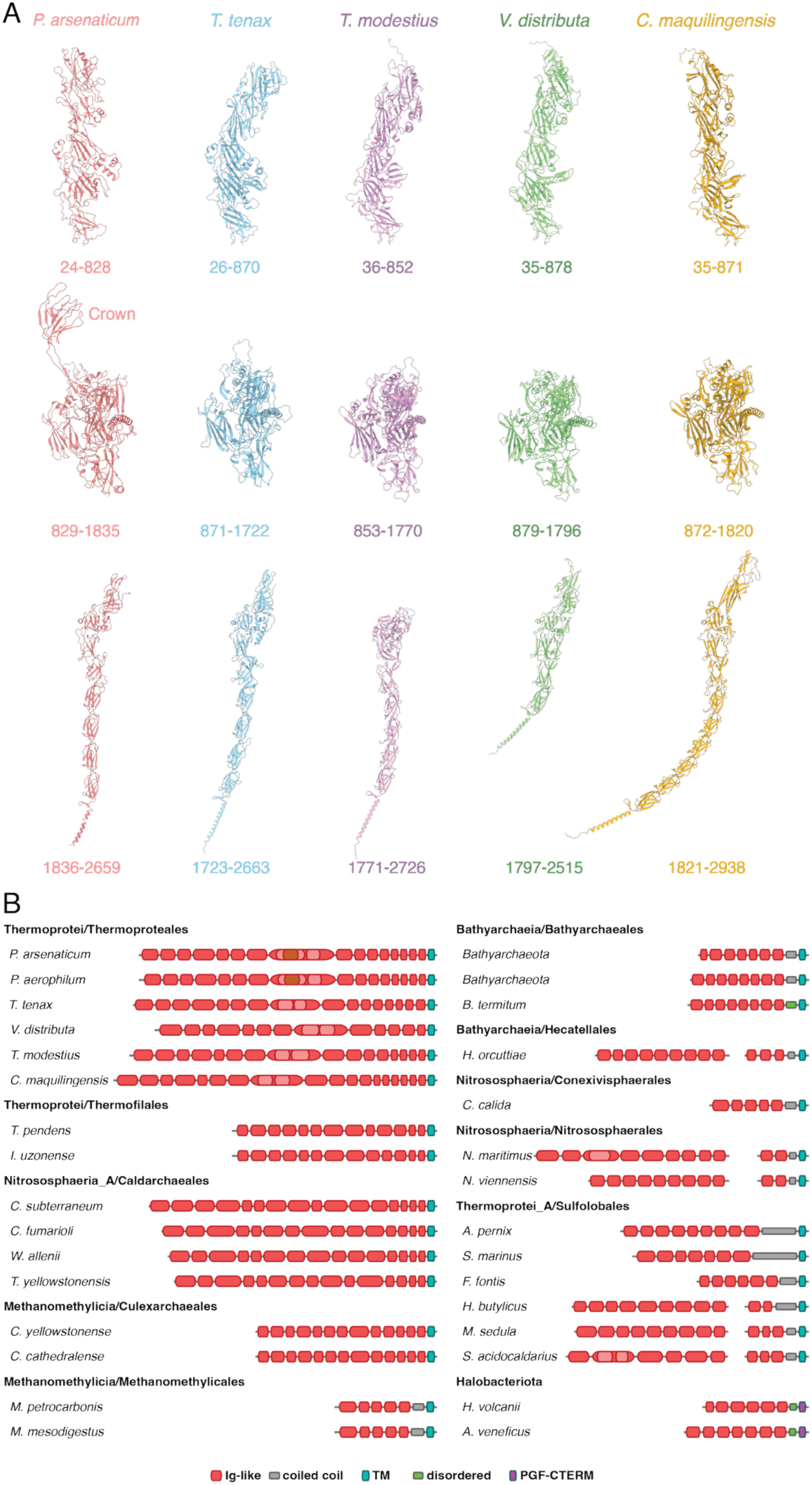
- AlphaFold3 predictions and predicted domain arrangements for candidate SLPs from other Thermoproteales. **A)** Comparison of the PySLP experimental model with AlphaFold3 predictions for the corresponding segments of *T. tenax* (UniProtKB G4RLQ9), *T. modestius* (UniProtKB A0A830GSK0), *V. distributa* (UniProtKB E1QP79) and *C. maquilingensis* (UniProtKB A8M8H4). The predicted structure generated for each segment was superposed onto PySLP and visualised from the same orientation, and residue range for each of the segments is noted. **B)** Predicted domain arrangement of the candidate SLPs identified by HHpred searches across Thermoproteota, together with two reference SLPs from Halobacteriota. Domains are shown according to their in their relative positions within each SLP and are coloured by their experimentally determined or predicted folds. Protein accession numbers are provided in Supplementary Table 2.

To gain further insight into the evolution of *Pyrobaculum* SLPs, we assembled a dataset of more closely related homologs from *Pyrobaculum* and *Thermoproteus* species, which allowed unambiguous alignment of the corresponding sequences. Mapping sequence conservation on the hexameric PySLP model revealed that the surface exposed to the extracellular environment displayed considerably higher divergence than the surface facing the cytoplasmic membrane (Supplementary Fig. 10A). A notable exception to this pattern was the Ig-like domains located at the trimer-stacking interface (Fig. 5B) and involved in S-layer stabilisation, which displayed higher sequence conservation (Supplementary Fig. 10A). Although we could not conclusively assign side chains, the PySLP sequence revealed an unexpectedly high number of cysteine (Cys) residues, all present within the lattice-forming region (Supplementary Fig. 10B). In total, each PySLP monomer contains 14 Cys residues, potentially forming up to seven putative disulfide bridges that could mediate both intradomain and interdomains interactions (Supplementary Fig. 10B), although these could not be confirmed at the resolution of our cryo-EM map. The Cys pairs potentially involved in disulfide bond formation were also conserved across *Pyrobaculum* SLPs, and most were also present in the *Thermoproteus* homologs (Supplementary Fig. 10C). Overall, this pattern of sequence conservation is consistent with diversification of the exposed S-layer surface in response to changing environmental conditions, viral infections, or competition with other microbes, while preserving regions important for lattice assembly and stability.

We next used HHpred^55^ to identify more divergent homologs of this family that may have escaped detection by BLAST. To this end, we searched the proteomes of representative species from the order Thermofilales within the class Thermoprotei, as well as from other classes of the phylum Thermoproteota. This analysis identified PySLP homologs in Methanomethylicia (order Culexarchaeales), including *Candidatus* Culexmicrobium cathedralense (GenBank MCS7385630.1) and *Ca*. Culexarchaeum yellowstonense (UniParc UPI0044657478); in Nitrososphaeria_A (order Caldarchaeales), including *Ca*. Caldarchaeum subterraneum (GenBank BAJ47616.1), *Ca*. Calditenuis fumarioli (GenBank MDJ0274653.1), *Ca*. Wolframiiraptor allenii (GenBank MCL7393743.1), and *Ca*. Terraquivivens yellowstonensis (GenBank MCL7394657.1); and in Thermoprotei (order Thermofilales), including *Thermofilum pendens* (UniProt A1RY64) and *Infirmifilum uzonense* (UniProt A0A0F7FI64). These share the overall PySLP-like architecture, including tandem Ig-like domains, the conserved membrane-proximal β-hairpin, the transmembrane helix, and the basic C-terminal tail. However, the homologs vary considerably in length and domain composition. The Culexarchaeales proteins are the most compact, while the Caldarchaeales proteins are substantially larger (Fig. 7B). Their increased size appears to result not only from a greater number of Ig-like domains but also from extensive insertions within several domains, which were largely absent from the more compact Methanomethylicia and Thermofilales homologs. Thus, although these divergent putative SLPs retain a conserved overall architecture and membrane-anchoring arrangement, they exhibit substantial lineage-specific variation through changes in Ig-like domain number and structural elaboration. We therefore propose that these proteins also form PySLP-like S-layers, but with lineage-specific adaptations. More broadly, we hypothesize that this conserved S-layer architecture contributes to cell-envelope stability in *Pyrobaculum* and other (hyper)thermophilic archaea and may represent a specialised adaptation to life at high temperatures.

S-layers in which the lattice is positioned at a considerable distance from the cytoplasmic membrane by an extended stalk are not unique to *Pyrobaculum*, but also occur in other Thermoproteota. In *Staphylothermus marinus*, for example, the *p*4 S-layer lattice is held approximately 70 nm above the membrane by membrane-anchored, tetrameric coiled-coil stalks^62^. In *Sulfolobus acidocaldarius*, the *p*3 SlaA lattice is positioned approximately 30 nm above the membrane by the membrane-anchored stalk protein SlaB, which comprises three Ig-like domains together with an extended coiled-coil region. S-layers with substantial space between the cytoplasmic membrane and the lattice may therefore represent a recurrent feature of Thermoproteota, although they are constructed using distinct molecular architectures. In *Pyrobaculum*, membrane anchoring, stalk formation, and lattice assembly are integrated within a single colossal Ig-rich protein, with the stalk itself formed by an array of Ig-like domains. By contrast, the *S. acidocaldarius* S-layer is a three-component system in which the Ig-rich lattice protein SlaA is supported by SlaB and Saci1846^7,29^, whereas the *S. marinu*s SLP combines a lattice-forming region and a long coiled-coil stalk within a single polypeptide. To investigate the possible evolutionary relationships among these systems, we performed all-against-all HMM-HMM (hidden Markov model) comparisons using HHpred on representative experimentally characterised and putative SLPs from across the classes of Thermoproteota, together with selected Halobacteriota reference SLPs. These comparisons revealed strong sequence similarity within each architectural group, together with extensive cross-group relationships throughout the phylum (Supplementary Fig. 11). In particular, PySLP-like proteins from Thermoprotei, Methanomethylicia, and Nitrososphaeria_A showed significant matches to one another, as well as to the Ig-like region of SlaB. Together, these findings suggest that the diverse S-layers of Thermoproteota may have evolved from related ancestral Ig-rich proteins through lineage-specific domain duplication, divergence, fusion, and structural elaboration in response to the ecological niche of the organisms.

## Discussion

S-layers are among the most widespread cell-envelope structures in prokaryotes, occurring across a variety of bacterial and archaeal phyla and often constituting the primary interface between the organism and its environment^6^. Despite their pervasiveness and importance for cell protection, shape determination, adhesion, environmental interactions, and cell-cell recognition^2^, the evolutionary origins of SLPs remain poorly understood^63^. Structures of SLPs determined from diverse organisms have revealed a remarkable diversity in protein architecture, ranging from relatively small proteins composed of only a few domains to exceptionally large multidomain proteins containing extensive arrays of repeating motifs or domains, as well as substantial differences in their oligomeric organisation^8,29,39,40,42,45,52,57,64^.

In archaea, there is evidence for divergent evolution among SLPs^63,65^. The evolution of large multidomain SLPs could have occurred through repeated domain duplication, amplification, and subsequent diversification^16,58^. Tandem arrays of structurally similar Ig-like domains are a recurring feature of archaeal S-layers^8,29,39,45^, and the extraordinary size and multidomain architecture of PySLP may represent an extreme example of this paradigm. Consistent with such an evolutionary scenario, the divergent PySLP-like proteins identified across Thermoproteota retain a similar overall architecture but vary considerably in the number of Ig-like domains and in the extent of structural elaborations within individual domains. Our HMM-HMM comparisons (Supplementary Fig. 11) further supported evolutionary relationships among several S-layer systems in Thermoproteota, with PySLP-like proteins from multiple classes showing significant similarity to one another and to SlaB-like proteins from Sulfolobus. Together, these observations suggest that the diverse S-layer systems of Thermoproteota may have evolved from related ancestral Ig-rich proteins through lineage-specific domain duplication, divergence, fusion, and structural elaboration. In particular, positioning the S-layer lattice far from the cytoplasmic membrane through an extended stalk, thereby creating a substantial “epiplasmic” space, may represent a recurrent architectural theme in Thermoproteota, realised through distinct single- and multi-component systems.

Beyond its unusual architecture and evolutionary relationships, PySLP also contains several features that may contribute to stability at high temperatures. As the first S-layer structure described from a hyperthermophilic archaeon, PySLP provides an opportunity to examine structural adaptations associated with life under extreme thermal conditions. In particular, PySLP contains multiple Cys residues that could form disulfide bonds, stabilising individual domains or interlock neighbouring domains within the lattice-forming region and crown domain (Supplementary Fig. 10B). Notably, many SLPs from mesophilic organisms are almost entirely devoid of Cys residues^40^. An increased number of disulfide bridges represents one of the adaptations to high-temperature environments in hyperthermophilic archaea and their viruses^66,67^, and has previously been observed in the *P. arsenaticum* type IV pilus^50^ and the *S. acidocaldarius* S-layer^29^. However, in both cases, only a single disulfide bond has been reported. PySLP therefore appears to rely more extensively on disulfide bonds for stabilisation. Furthermore, it oligomerises into a highly interconnected hexamer stabilised by extensive interactions within the pillar and crown regions. It also displays an overall highly charged surface (Supplementary Fig. 8B) and contains multiple putatively assigned N-glycosylation sites distributed across the S-layer (Supplementary Fig. 8A).

The precise physiological role of the colossal S-layer found in *P. arsenaticum* and other members of the *Pyrobaculum* genus remains uncertain. However, the considerable energetic investment required for its production and the large number of copies needed to coat an entire cell suggest that it performs essential biological functions. One likely function is the maintenance of cell morphology and envelope integrity. Cryo-ET revealed highly rigid, rod-shaped cells, supporting the proposal that archaeal S-layers function as exoskeleton-like structures that are critical for cell-shape determination and mechanical stability, particularly in organisms lacking a peptidoglycan cell wall^68^. The stability of the cell shape may be further reinforced by anchoring of the S-layer to the cytoplasmic membrane, which can modify the membrane’s biophysical properties, reduce its elasticity, and help maintain membrane integrity at elevated temperatures^10^.

Beyond this structural role, the unusual architecture of the *Pyrobaculum* S-layer may support additional functions at the cell surface. Notably, in *Pyrobaculum* and *Thermoproteus* species, the SLP gene is found within a conserved gene neighbourhood. The potential operon includes genes encoding a membrane-associated Ig-like domain-containing protein, a functionally uncharacterised AMMECR1-domain protein, a thioredoxin-like fold-containing protein, an adenylate cyclase, and an ATP-dependent DNA ligase (Supplementary Fig. 12). Among these, the presence of the DNA ligase is particularly intriguing and raises the possibility of a functional relationship between the cell surface and extracellular DNA. Given the highly positively charged surface of the S-layer and the approximately 12-Å-wide channel formed through the centre of the pillar, one possibility is that the S-layer facilitates the uptake of extracellular DNA from the surrounding environment. Such a function could be particularly relevant during biofilm formation, as *Pyrobaculum* species also encode AbpA/AbpB proteins implicated in archaeal biofilm matrix assembly^49^.

More broadly, this exceptional S-layer architecture may represent an adaptation to life in hyperthermal environments. We identified closely related, uncharacterised SLP candidates in several thermophilic and hyperthermophilic members of the order *Thermoproteales*, including *T. tenax*, *T. modestius*, *V. distributa*, and *C. maquilingensis* (Fig. 7). More divergent candidates with the same overall architecture were also identified in the orders Thermofilales, Caldarchaeales, and Culexarchaeales, extending its distribution across multiple classes of Thermoproteota. These findings suggest that this architecture is conserved across diverse thermophilic and hyperthermophilic lineages inhabiting similar environmental conditions. The evolution of an exceptionally large, highly interconnected protein lattice may therefore provide an additional layer of structural reinforcement that helps maintain cellular integrity under these conditions. The S-layer is also likely to represent a major interface with viruses infecting these organisms. It will therefore be important to investigate the interplay between the protection afforded by these colossal S-layers and the evolution of the viruses themselves. In particular, it will be interesting to understand how viruses of *Thermoproteales*^69^ overcome this formidable barrier and deliver their genetic material into the archaeal cell. Future genetic and biophysical studies will be needed to determine whether these colossal S-layers confer measurable advantages in thermotolerance or cellular protection and how their presence represents a specific adaptation to life at higher temperatures.

## Methods

### Growth of P. arsenaticum and P. oguniense cells

Liquid culture of *P. arsenaticum* strain 2GA^33^ was grown in DSMZ medium 1090 at 90 °C without shaking. *P. oguniense* TE7 (DSM 13380) was purchased from the DSMZ and grown in DSMZ medium 390 at 90 °C without shaking^70^.

### *P. arsenaticum* cell envelope purification

Cells from 50 ml of culture were collected by centrifugation (Eppendorf S-4xUniversal rotor, 4,347×g, 15 min, 20 °C), resuspended in 40 ml of buffer (10 mM NaCl, 1 mM phenylmethylsulfonyl fluoride, 0.5% (w/v) sodium lauroylsarcosine, and 10 μg/ml DNase I) and incubated in a shaker with agitation (140 rpm) at 37 °C for 2 hours. Then, samples were pelleted by centrifugation (Beckman JA-17 rotor, 18,000×g, 30 minutes, 15 °C). The procedure was repeated twice, the pellet resuspended in 40 ml of the buffer (10 mM NaCl, 0.5% (w/v) sodium lauroylsarcosine) and incubated in a shaker with agitation (140 rpm) at 37 °C, overnight. Following centrifugation (as above), the resultant pellet was resuspended in 1.5 ml buffer (10 mM NaCl, 0.5 mM MgSO4, 0.5% (w/v) SDS) and incubated in a shaker with agitation (140 rpm) for 20 min at 37 °C. Purified S-layer sacculi were washed four times with distilled water and stored at 4 °C until further experimentation.

### Sample preparation for cryo-EM and cryo-ET

For cryo-EM SPA of cell envelopes, 2.5 μl of the purified S-layer sample was applied to glow discharged Quantifoil R2/2 Cu/Rh 200 mesh grids, incubated for 2 seconds and blotted for 5 seconds before plunge freezing into liquid ethane using a Vitrobot Mark IV (Thermo Fisher Scientific) at 10 °C and 100% humidity. For cryo-ET, 2.5 μl of *P. arsenaticum* 2GA or *P. oguniense* cells, mixed with 10 nm gold particles conjugated to protein A, were applied on glow discharged Quantifoil R3.5/1 Cu/Rh 200 mesh grids, incubated for 10 seconds and blotted for 3.5 or 3 seconds before plunge freezing into liquid ethane using a Vitrobot Mark IV at 10 °C and 100% humidity.

### Cryo-EM and cryo-ET data collection

Single particle cryo-EM data was collected using a Titan Krios G3 microscope (Thermo Fisher Scientific) running at 300 kV equipped with a Quantum energy filter (slit width 20 eV) and a K3 direct electron detector (Gatan) at 81,000 nominal magnification and a pixel size of 0.546 Å running in counting super-resolution mode. A total of 2,530 movies of the untilted specimen and 4,096 movies of 30° tilted specimen were collected using EPU (Thermo Fisher Scientific), with a total dose of 49.6 or 52.2 e^-^/Å^2^, respectively, with each movie consisting of 40 frames.

Cellular cryo-ET data of *P. arsenaticum* was collected using a Titan Krios G4 microscope (Thermo Fisher Scientific) running at 300 kV equipped with a SelectrisX imaging filter (slit width 10 eV) and a Falcon 4i direct electron detector (Thermo Fisher Scientific) at 26,000 nominal magnification and a pixel size of 4.71 Å using the SerialEM program^71^. Each tilt-series was acquired with a total dose of 121 e^-^/Å^2^ applied over the complete series and split equally over the tilt range of ±60° with 1° tilt increments. 22 tilt-series were acquired at a defocus range of -3 to -7 μm and an exposure time of 2.622 seconds per tilt image.

Cellular *in situ* data of *P. oguniense* was collected using a Titan Krios G3 microscope (Thermo Fisher Scientific) running at 300 kV equipped with a Gatan imaging filter (slit width 20 eV) and a K2 direct electron detector (Gatan) at 26,000x magnification and a pixel size of 5.527 Å using the SerialEM program. Each tilt-series was acquired with a total dose of 140 e^-^/Å^2^ applied over the complete series and split equally over the tilt range of ±60° with 1° tilt increments. Four tilt-series were acquired at a defocus range of -6 to -8 μm and an exposure time of 3 seconds per tilt image, dose-fractioned across 10 frames.

### *P. arsenaticum* S-layer SPA data processing

Movies from untilted (2,530 movies) and 30° tilted (4,096 movies) data collections were initially processed independently using RELION4.0^72^. Movies were clustered into optics groups using a k-means algorithm (https://github.com/DustinMorado/EPU_group_AFIS). Imported movies were motion corrected, dose-weighted and Fourier cropped with MotionCor2 implemented in RELION4.0^72,73^ and CTF estimation of motion-corrected micrographs was performed using CTFFIND4^74^. Using the helical picking tool in RELION4.0, top and side views of the S-layer lattice were manually picked, downsampled by a factor of 4 to a box of 128 X 128 pixels^2^, and these were subjected to two-dimensional classification in RELION4.0. Using the coordinates of particles from good class averages as a training dataset, particles were picked using TOPAZ^75^, classified in two dimensions, resulting in 296,237 particles. These were re-extracted to 512X512 pixels^2^ box at 1.092 Å/pixel, subjected to three-dimensional refinement in RELION4.0 with C6 symmetry applied, which generated a 6.7 Å resolution reconstruction using the gold-standard 0.143 FSC criteria (Supplementary Fig. 2).

Further processing was performed in CryoSPARC^76^, starting with patch CTF estimation, then using the previous model as a template, 7,708,473 particles were picked using the CryoSPARC template picker. These were downsampled by a factor of 3 and extracted to a 220X220 pixel^2^ box, then subjected to two-dimensional classification rounds, and 500,269 particles were retained for subsequent processing. This particle subset was downsampled (from the original unbinned pixel) by a factor of 2 and extracted into a 282X282 pixel^2^ box. Next, duplicate particles were removed and then were subjected to homogeneous refinement, as well as non-uniform refinement^77^ resulting in a 5.5 Å resolution reconstruction when applying C6 symmetry, with the pillar poorly resolved. This was followed by multiple 3D classification rounds with C1 symmetry, and these resulted in a subset of 67,152 particles extracted at 440X440 pixel^2^ box at a 1.5 downsampling factor with a final pixel size of 1.638 Å. These were subjected to non-uniform refinement using C6 symmetry, complemented with local CTF refinement^78^ in CryoSPARC, which resulted in a final 5.75 Å resolution reconstruction using the gold-standard 0.143 FSC criteria (Supplementary Figs. 2-3), with local resolution as low as 4.8 Å (Supplementary Fig. 3G).

### *P. arsenaticum* cell envelope peptide fingerprinting mass-spectrometry

The protein content of the *P. arsenaticum* S-layer preparations was analysed by liquid chromatography – tandem mass spectrometry (LC-MS/MS) at the Proteomics Platform of Institut Pasteur (Paris, France) as previously described^79^. Peptide masses were searched against the UniProt proteomes of *P. arsenaticum* strains DSM 13514 (UP000001567) and 2GA (UP000554766), respectively using Andromeda^80^ with the MaxQuant ver. 2.0.3.0 software^81^. Variable modifications (methionine oxidation and N-terminal acetylation) and fixed modification (cysteine carbamidomethylation) were set for the search, and trypsin with a maximum of two missed cleavages was chosen for searching. The minimum peptide length was set to 7 amino acids, and the false discovery rate (FDR) for peptide and protein identification was set to 0.01. The main search peptide tolerance was set to 4.5 ppm and to 20 ppm for the MS/MS match tolerance.

### PySLP structure prediction and model fitting into the cryo-EM map

The protein sequence from UniProt entry A0A7L4PAH2 was submitted to AlphaFold3 server^37,44^, however the predicted structure could not be fitted confidently to the cryo-EM SPA map. To improve the predictions, the protein sequence was divided to three roughly sized segments - residues 24-828, 829-1835 and 1836-2659. The final 8 residues (2660-2667) were trimmed from the final structure due to poor map coverage. The crown domain (residues 1312-1461) was predicted separately as six copies as they are present at the top centre of the pore-like pillar density. The individual segments were manually fit the cryo-EM map using Coot^82^, keeping all secondary structures rigid, and merged together to generate a single chain. The single chain model was subsequently refined using Servalcat^83^ and PHENIX^84^, then duplicated to form a C6 symmetrical model, and refined using the cryo-EM map. Figure panels were prepared using ChimeraX^85^.

### Cryo-ET data processing and S-layer *in situ* STA

*P. oguniense* tilt-series were processed using the IMOD package^86^ and aligned using gold fiducials. To improve visualisation the tomograms were produced using the simultaneous iterative reconstruction technique (SIRT) implemented in Tomo3D^87,88^.

*P. arsenaticum* tilt-series were processed using RELION5.0 software^89^ as follows. Frames were imported and motion corrected using the RELION MotionCor2 implementation^73^, and CTF estimation was performed using CTFFIND4^74^. Tilt-series were aligned utilising gold fiducials using the IMOD^86^ RELION5.0 wrapper and the fiducials were removed from the final tomograms using FIDDER (https://github.com/teamtomo/fidder). Tomograms were reconstructed and CTF corrected using RELION5.0 and then denoised with Cryo-CARE^90^.

Using the cryo-EM SPA map as a template, S-layer particles were picked within the non-denoised tomograms using PyTOM template matching^91^. In total, 20,273 subtomograms were selected with their Euler angles assigned by PyTOM. These were subjected to three-dimensional classification in RELION5.0 to clean up the dataset combined with two iterations of extending the lattice positions using a custom script described previously^8^ followed by three-dimensional classification, and these resulted in a total of 8,429 2D stacks that were extracted to an unbinned 144X144 pixels^2^ box. These 2D stacks were then refined in three dimensions to produce an 18 Å-resolution reconstruction using the gold-standard 0.143 FSC criteria (Supplementary Fig. 13). To resolve the lattice arrangement, this subset of 2D stacks was extracted to a 256X256 pixel^2^ box downsampled to a factor of 4 (final box size 64X64) and 3D refined using RELION5.0. Data visualisation was prepared using ChimeraX^85^, ArtiaX^92^, and IMOD^86^.

### Bioinformatic analyses of the *P. arsenaticum* S-layer

The sequence of PySLP (UniProtKB A0A7L4PAH2) was used in protein-protein BLAST^59^ against the RefSeq^93^ and ClusteredNR^94^ databases. These searches identified many sequences belonging to members of the *Pyrobaculum* genus, as well as several uncharacterised, large protein sequences (over 2000 amino acids) with 23-39% sequence identity with PySLP. These belong to the Thermoproteaceae family and include *T. tenax*, *T. modestius*, *V. distributa*, and *C. maquilingensis*. To identify remote homologs of PySLP and other experimentally characterised SLPs from the phylum Thermoproteota, we performed HHpred searches against the proteomes of representative Thermoproteota species^55^. In addition to PySLP, experimentally characterised SLPs used as queries included those from *Nitrosopumilus maritimus* (SlaA, UniProtKB A9A4Y9; SlaB, A9A4Y8), *Aeropyrum pernix* (Q9YEG7), *Staphylothermus marinus* (Q54436), and *Sulfolobus acidocaldarius* (SlaA, Q4J6E5; SlaB, Q4J6E6). Candidate SLPs for further analyses were selected based on homologs identified in these searches, together with experimentally characterised and putative SLPs reported in previous studies^63^.

For systematic comparison of sequence relationships among SLPs within Thermoproteota, we selected a total of 48 representative proteins, comprising SLPs from across Thermoproteota together with selected Halobacteriota reference SLPs (see Supplementary Table 2). Representative proteins were chosen to capture the major SLP architectures and taxonomic groups across Thermoproteota. Signal peptides, transmembrane helices, and coiled-coil segments were excluded from the sequences used for the comparison, retaining the Ig-domain-containing segments. Multiple sequence alignments were generated for each protein using HHblits^95^ against the UniRef30 database^96^ with two iterations and otherwise default parameters. The resulting A3M alignments were compared directly in an all-against-all manner using HHalign from the HH-suite3 package^95^. Pairwise E-values were transformed to −log_10_ (E-value) and capped at 10 for visualisation, such that matches with E-values ≤10^-10^ were displayed at the maximum intensity. The resulting similarity matrix was visualised as a heatmap using Python and Matplotlib. DeepTMHMM^46^ was used to predict signal peptides and transmembrane helices. Protein structures were predicted using the AlphaFold3 server^44^ or the ColabFold implementation of AlphaFold2^38^.

## Data availability

The cryo-EM map produced in this study has been deposited at the Electron Microscopy Data Bank (EMDB) under accession code EMD-59407. The atomic coordinates of the PySLP hexamer have been deposited in the RCSB Protein Data Bank under accession code 33FC.

## Acknowledgements

T.A.M.B. acknowledges the support of the Medical Research Council, as part of UK Research and Innovation (also known as UK Research and Innovation) (programme MC_UP_1201/31 to T.A.M.B.). T.A.M.B. and I.C. thank the Human Frontier Science Program (grant RGY0074/2021), the European Molecular Biology Organization, the Wellcome Trust (grant 225317/Z/22/Z), the Leverhulme Trust, and the Lister Institute for Preventative Medicine for support. I.C. was supported by an EMBO Long-Term Fellowship (ALTF 92-2022). The authors acknowledge Jonathan Doye (University of Oxford) for initial assistance in the AlphaFold prediction and protein assignment.

## Contributions

T.A.M.B. and M.K. conceived and supervised the project. V.C. and M.K. performed cell growth, S-layer purification and mass spectrometry analyses. I.C., S.v.D., A.v.K., Z.F. and T.A.M.B performed cryo-EM and cryo-ET studies. I.C. and V.A. performed the bioinformatic analyses. I.C. wrote the initial draft of the manuscript. S.v.D., A.v.K., Z.F., V.A., M.K. and T.A.M.B. revised and edited the manuscript.

---

**Supplementary Fig. 1.**
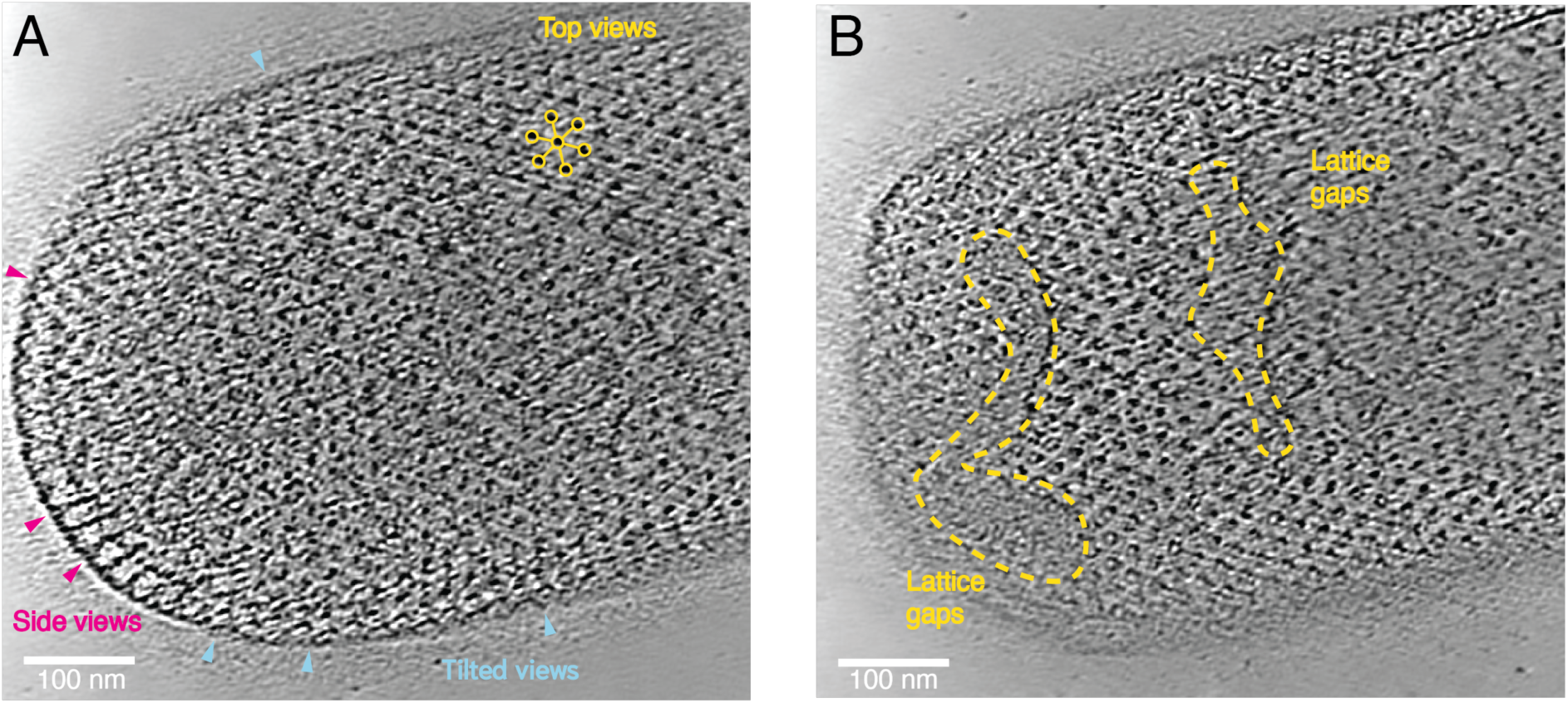
- Gaps in the *P. arsenaticum* S-layer lattice seen in cryo-ET. **A)** Tomographic slice of a *P. arsenaticum* cell showing the S-layer from top (yellow circles show the centre of individual hexamers), tilted (cyan arrowheads), and side views (magenta arrowheads). **B**) Tomographic slice from the same cell at a different height, displaying missing hexamers in the S-layer lattice. The gaps are marked with a dotted yellow line.

**Supplementary Fig. 2.**
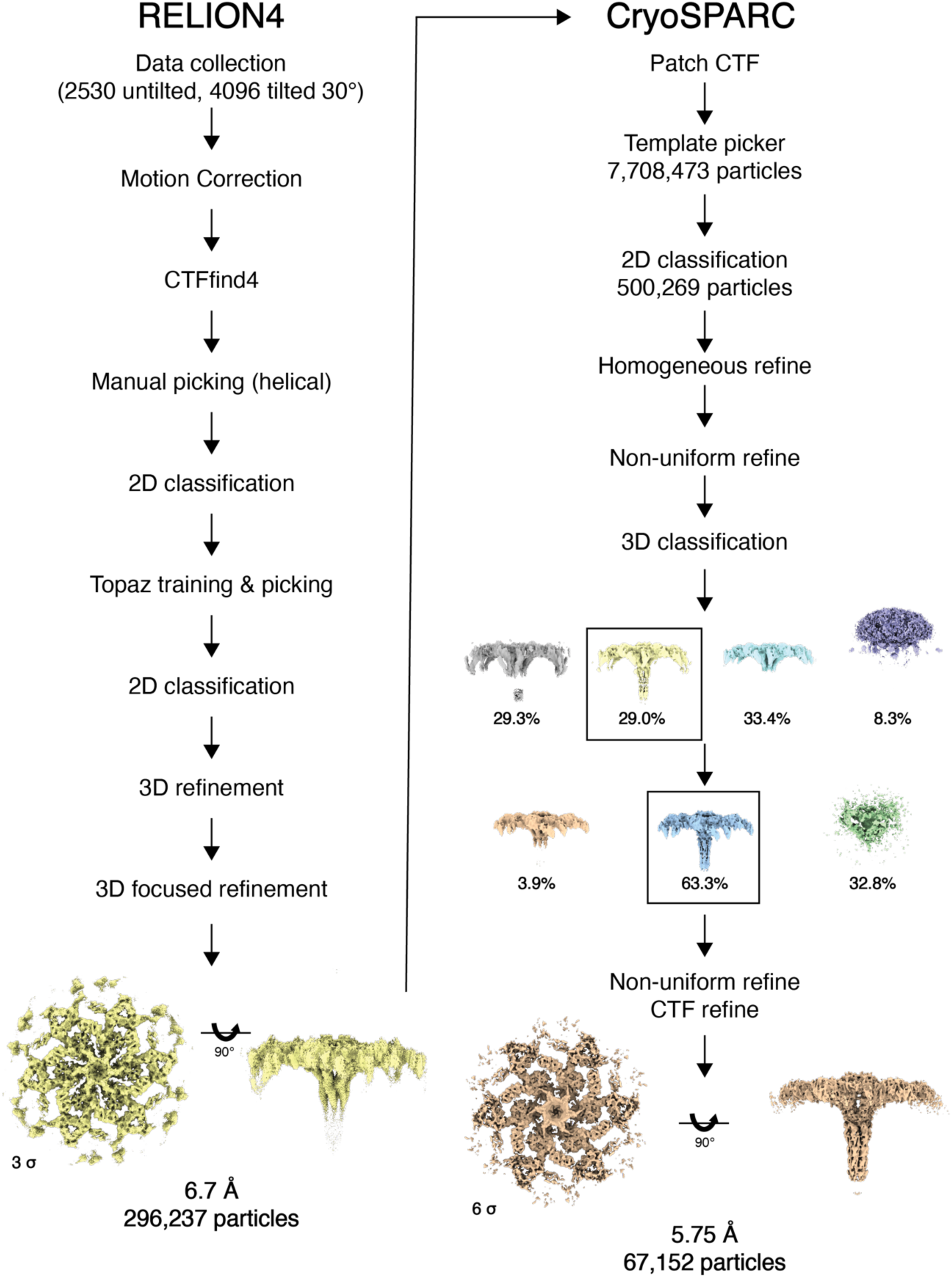
- Schematic of cryo-EM SPA data processing pipeline used. Left - data processing using the RELION4 workflow, resulting in a 6.7 Å reconstruction. The data was imported to CryoSPARC, and the RELION4 map was used as a template for further processing in CryoSPARC, which concluded with a final reconstruction at 5.75 Å resolution.

**Supplementary Fig. 3.**
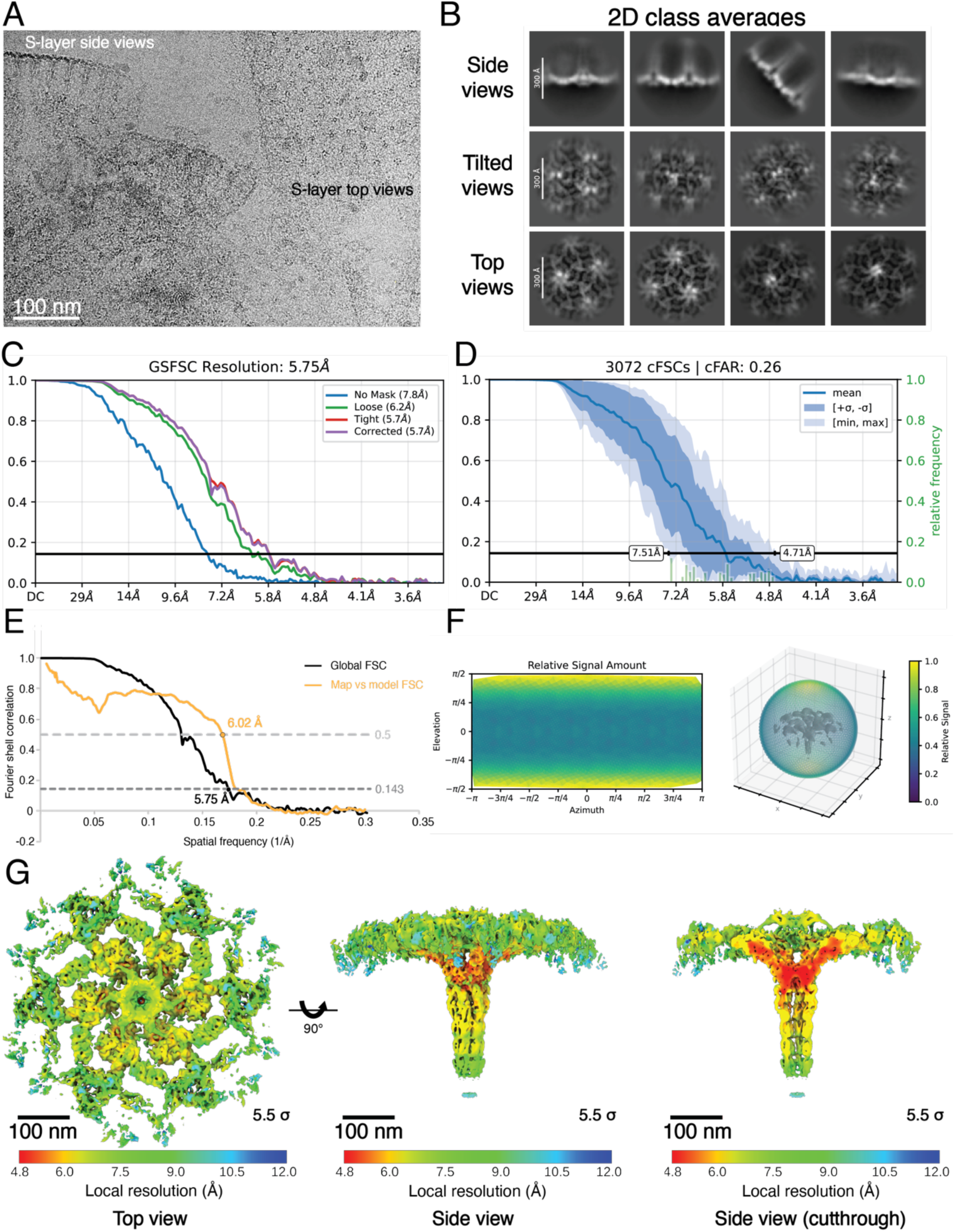
- Cryo-EM SPA resolution estimation. **A)** Cryo-EM image of purified S-layer sheets showing top and side views. **B**) Two-dimensional class averages from various orientations. **C**) Fourier shell correlation (FSC) resolution estimation of the cryo-EM SPA map. **D**) Conical FSC (cFSC) analysis of the two half maps. **E**) Map global FSC (black curve) and map-vs-model FSC curve (orange). **F**) Particle orientation analysis and their relative contribution to the reconstructed cryo-EM map. While top and bottom views dominate the reconstruction, tilted and side views are also present in the data set. **G**) Local resolution estimation showing the cryo-EM map from a top (left) and side view (centre), as well as a side view with slice through (right) showing that the local resolution is highest in the middle of the core hexamer.

**Supplementary Fig. 4.**
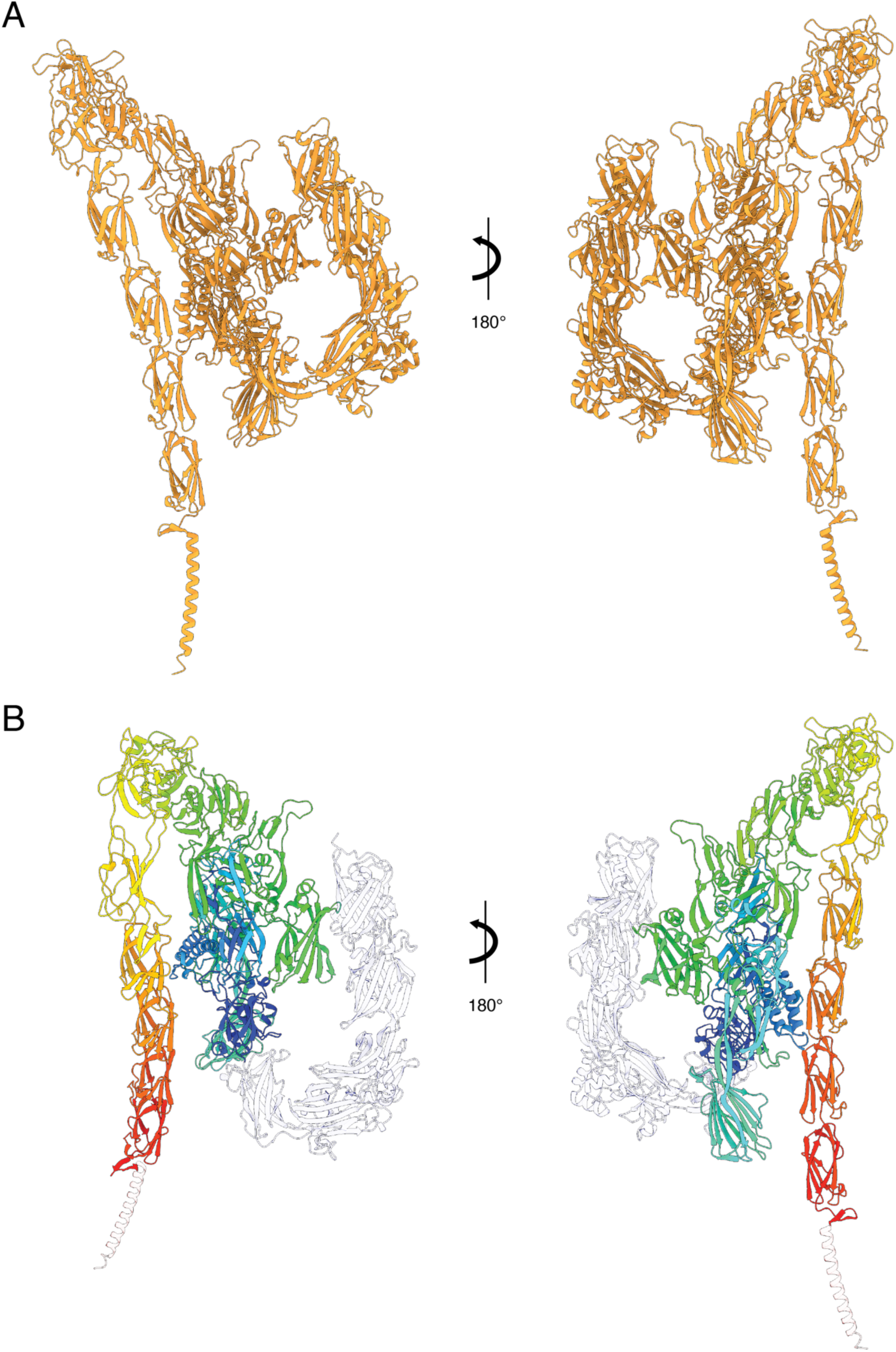
- AlphaFold3 prediction of the full length PySLP. **A**) The prediction differs significantly from the overall arrangement of the SLP observed in our cryo-EM SPA map. **B**) Prediction coloured by domains as in Fig. 3C.

**Supplementary Fig. 5.**
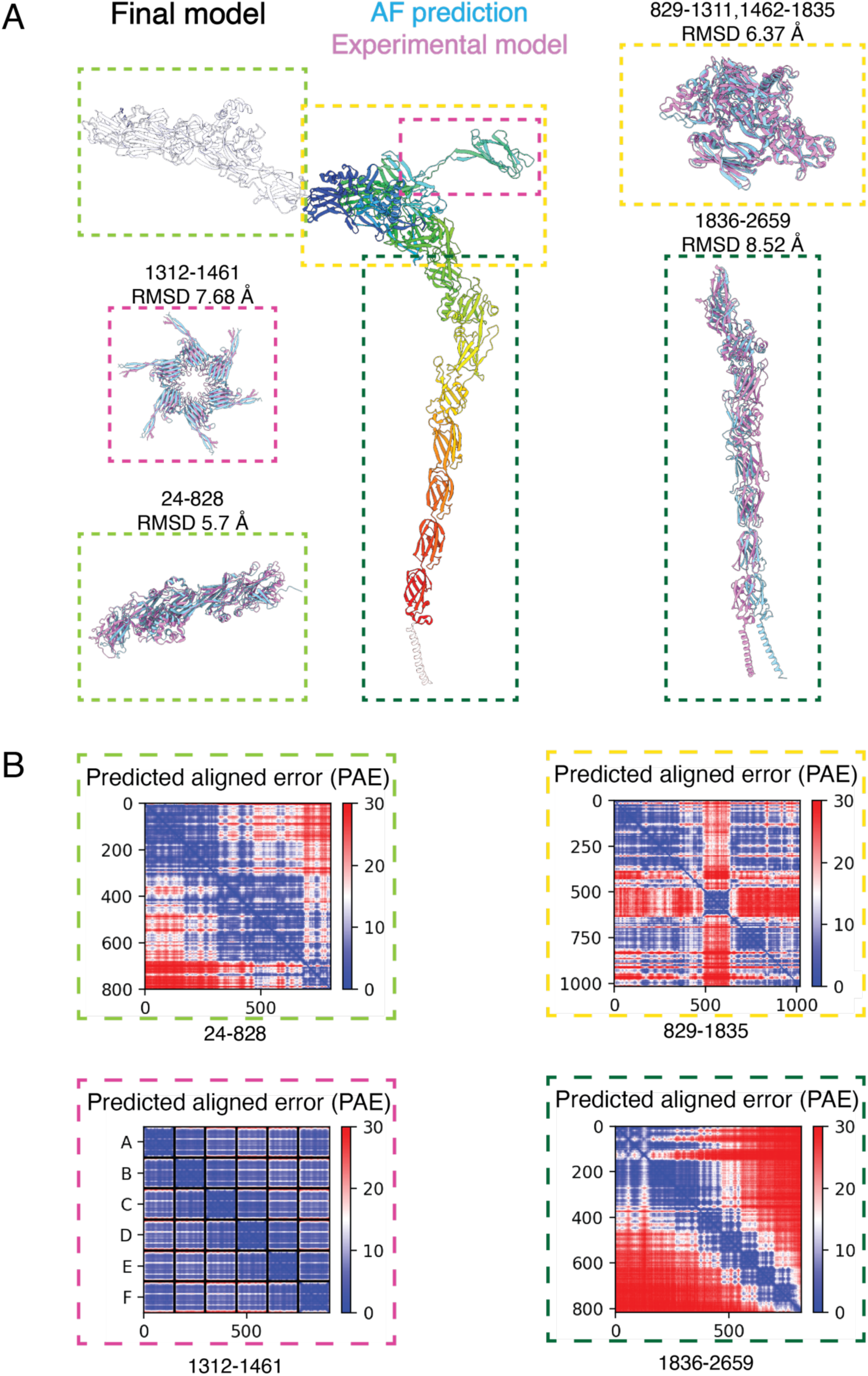
- AlphaFold prediction quality of the PySLP segments. **A**) Final model of PySLP monomer (coloured as in Fig. 3C) with the different segments for AlphaFold prediction marked by dotted rectangles - residues 24-828 (green), 829-1311 and 1462-1835 (yellow), 1312-1461 (magenta), and 1835-2659 (dark green). Each rectangle shows a superposition of the predicted and experimental models coloured cyan and plum, respectively, together with the RMSD values for each pair. Residues 1312-1461 (crown domain) were predicted as six copies (i.e. as a hexamer) to generate the hexameric centre. **B**) Predicted aligned error plots for each of the segments displaying per-residue AlphaFold confidence of their relative position, used to assess relative domain positioning. The error is measured in Å.

**Supplementary Fig. 6.**
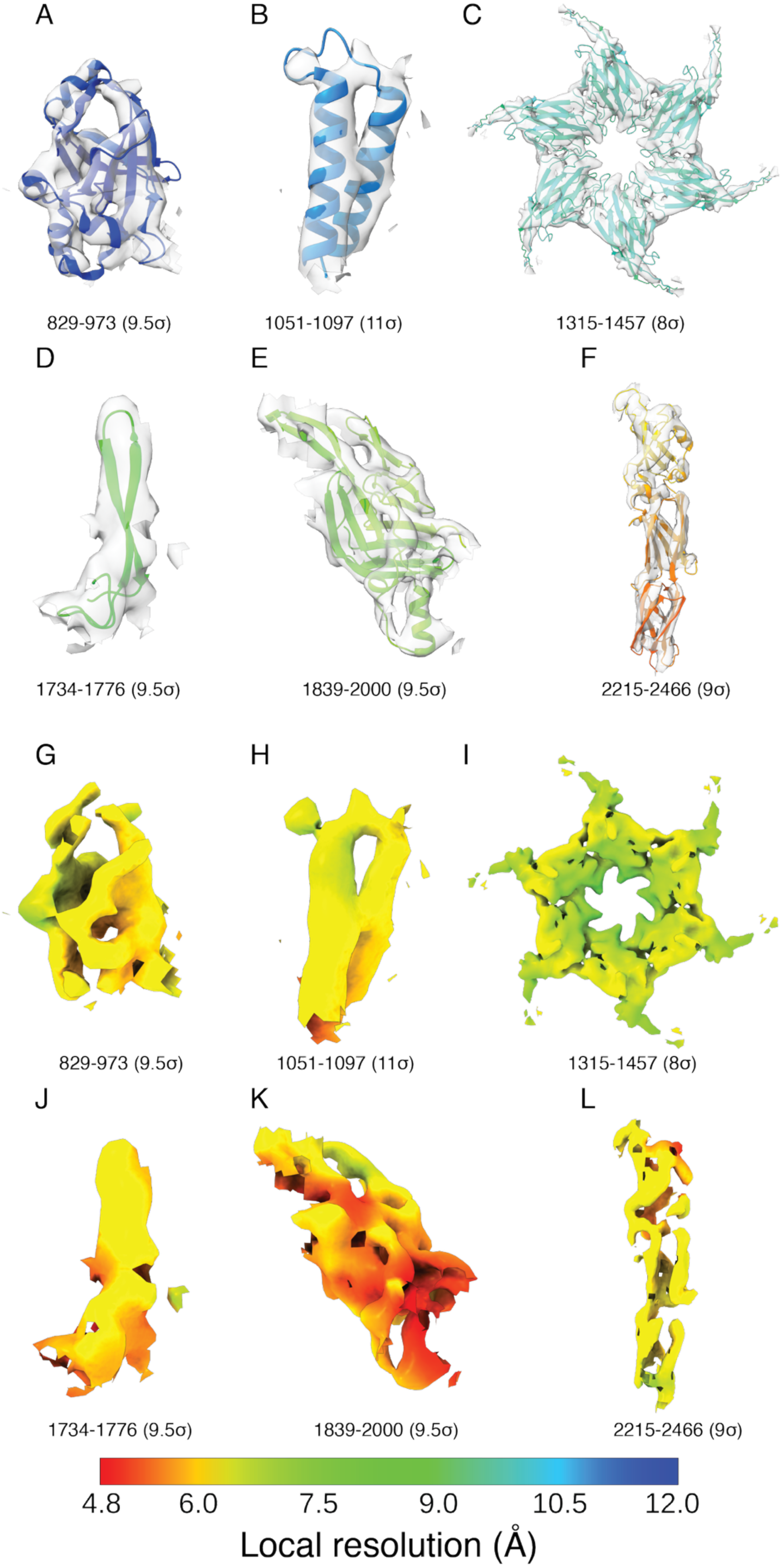
- Secondary structural elements and domains resolved in the cryo-EM map. **A-F**) Secondary structural elements or entire domains resolved in the cryo-EM map. The model is coloured as in Fig. 3C, the residue range and map contour level are marked, and the cryo-EM map is coloured in transparent light grey. **G-L**) Cryo-EM map of the same regions as in panels A-F coloured by local resolution.

**Supplementary Fig. 7.**
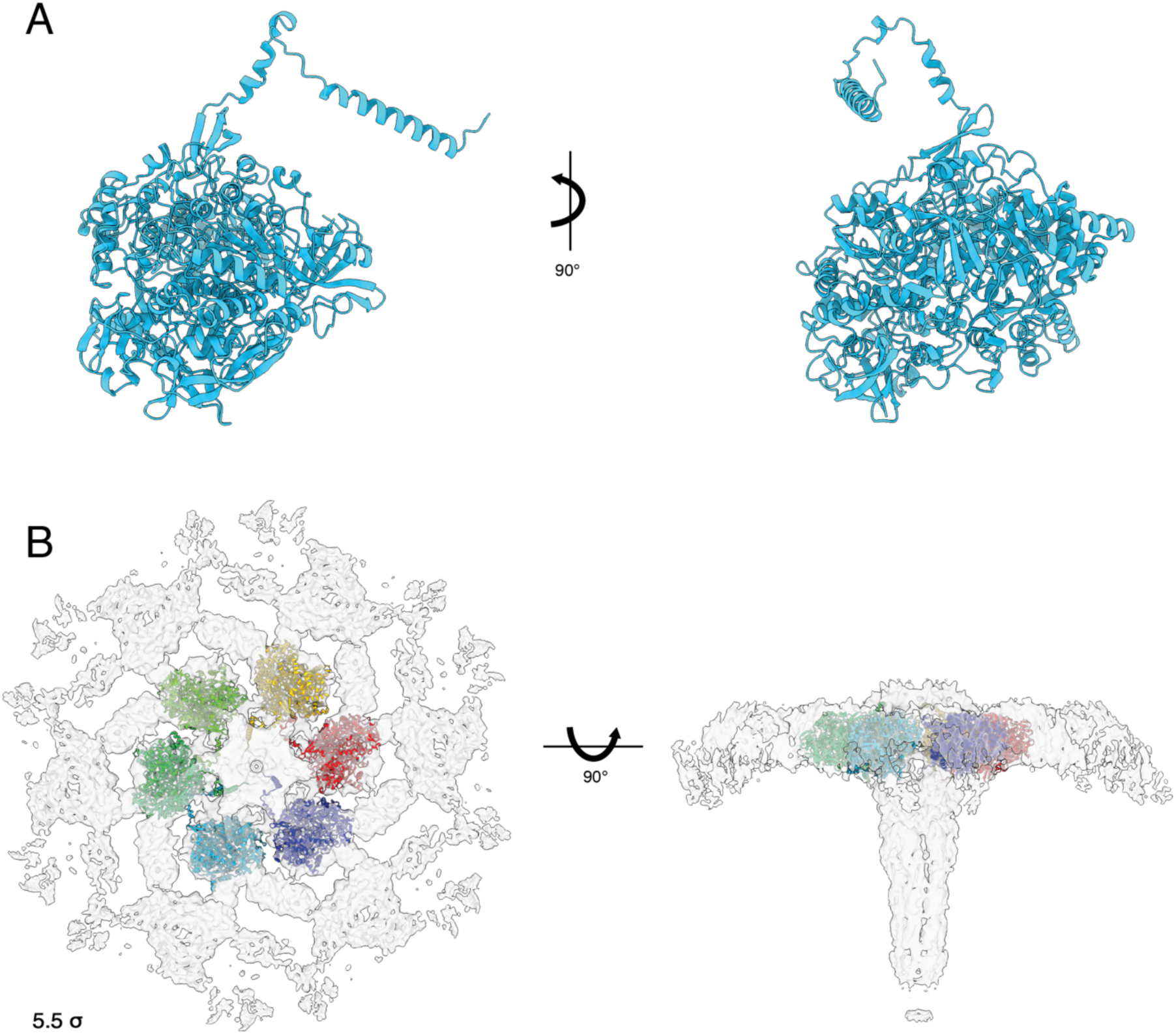
- AlphaFold prediction of the next best SLP candidate identified in peptide fingerprinting MS. **A**) Predicted model of another top hit in the peptide fingerprinting MS data (UniProtKB A0A7L4P773), annotated as a nitrate reductase. **B**) Six copies of the predicted structure fitted into the cryo-EM map. Each monomer is coloured separately, and the map is shown in transparent light grey, and the map contour level is marked. The model cannot explain all density observed in the map.

**Supplementary Fig. 8.**
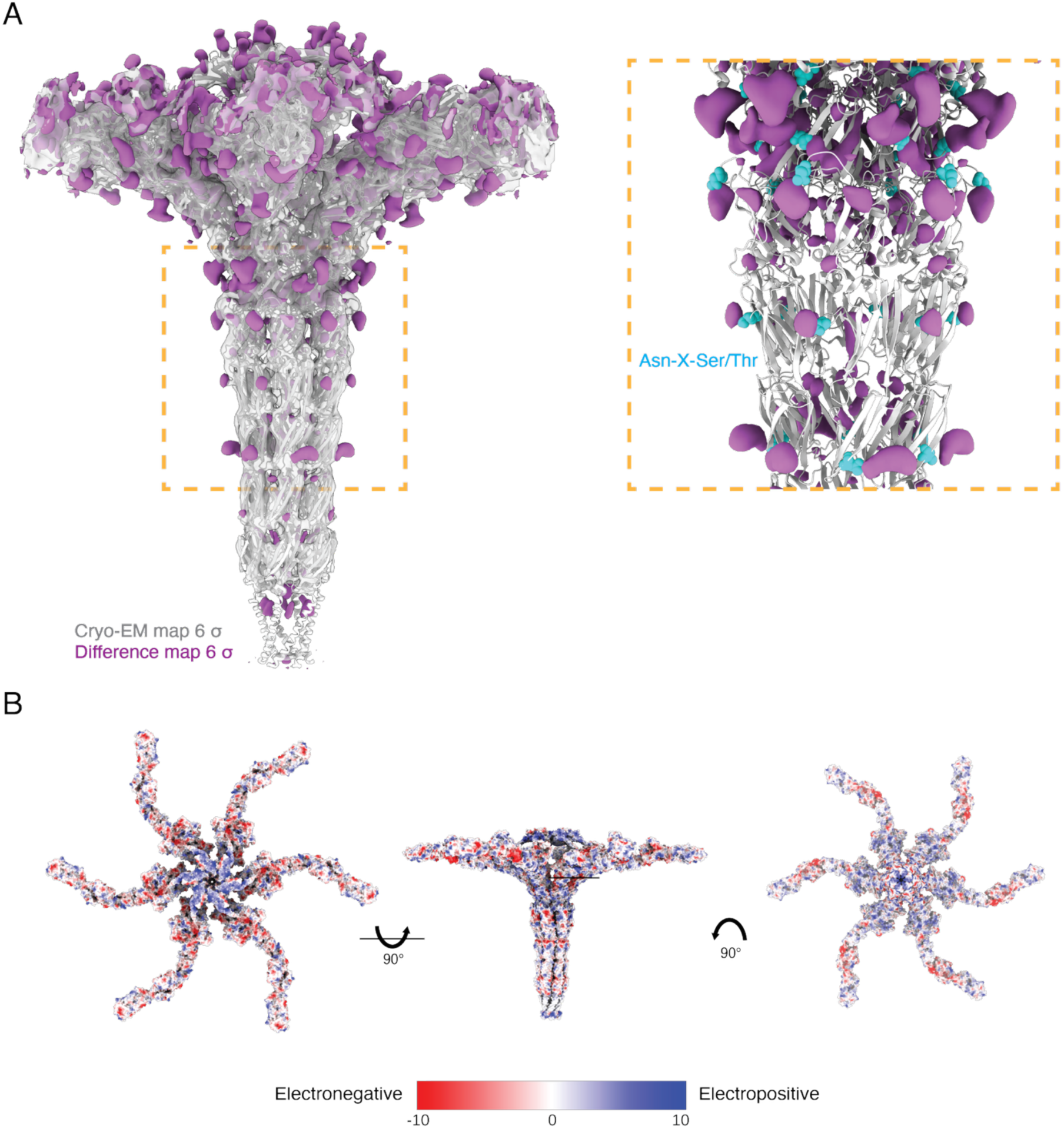
- Cryo-EM densities that could putatively be N-glycosylation sites and surface electrostatic potential of PySLP. **A**) Overlay of the cryo-EM SPA map and the difference map with the atomic model generated by Servalcat^97^ exhibiting multiple unassigned densities on the model surface. Rectangle shows the densities in the pillar C-terminal region. The protein is displayed as white ribbons, cryo-EM map in transparent light grey, and difference map in purple. Asparagine residues that correspond to the N-glycosylation motif (Asn-X-Ser/Thr) are shown as cyan spheres. Cryo-EM map and difference map contour levels are marked. **B**) Surface representation of PySLP hexamer coloured by its coulombic electrostatic potential. Electronegative regions are coloured red, electroneutral in white and electropositive in blue. The PySLP hexamer is shown from outside the cell (left), side view (middle) and from the cell looking outwards (right).

**Supplementary Fig. 9.**
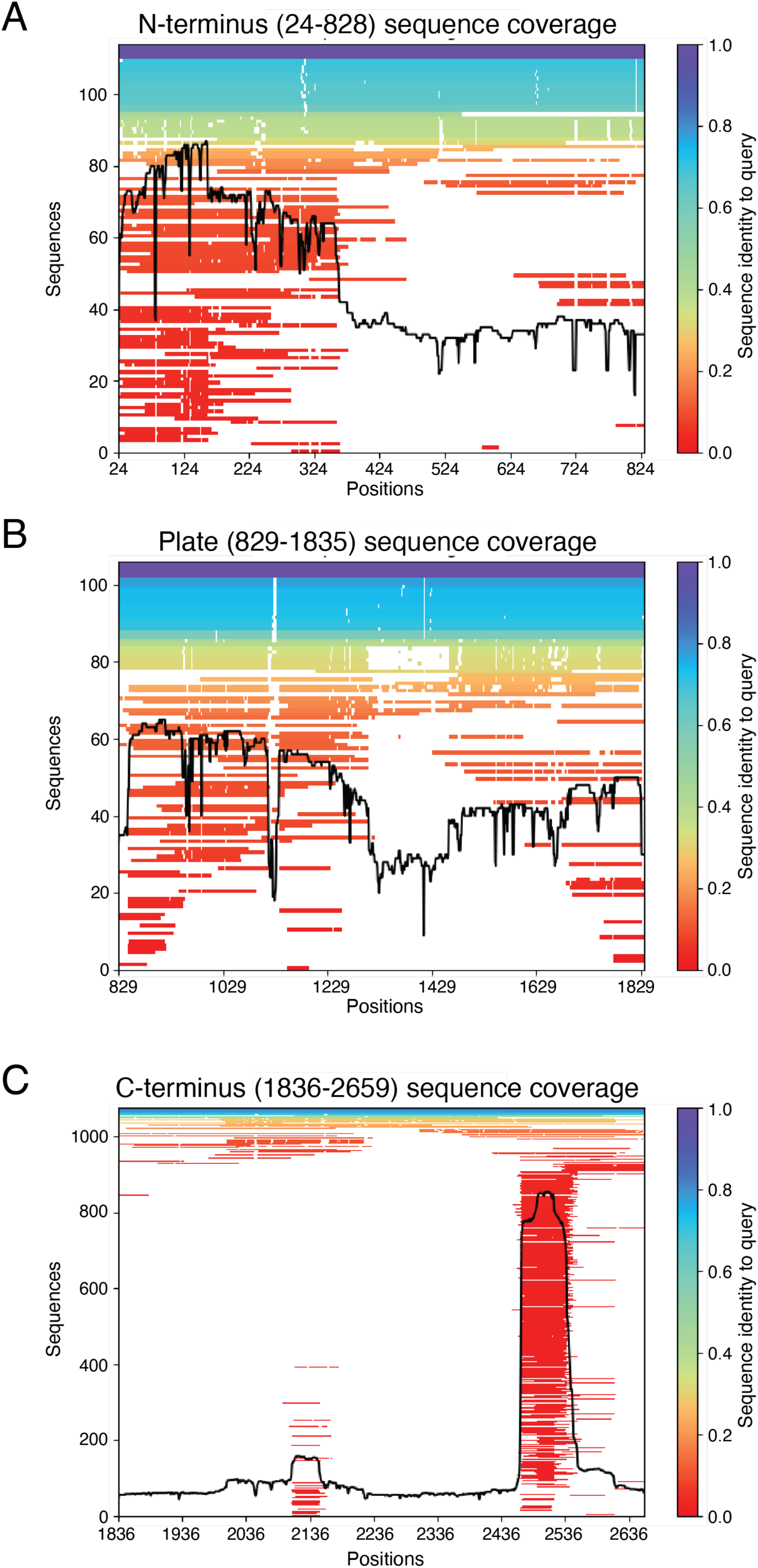
- ColabFold sequence coverage for predicted PySLP segments. Number of sequences identified by Colabfold^38^ multiple sequence alignment and their relative sequence identity to PySLP segments of the N-terminal region (**A**), central plate region (**B**) and C-terminal region (**C**). The residue range is noted for each segment.

**Supplementary Fig. 10.**
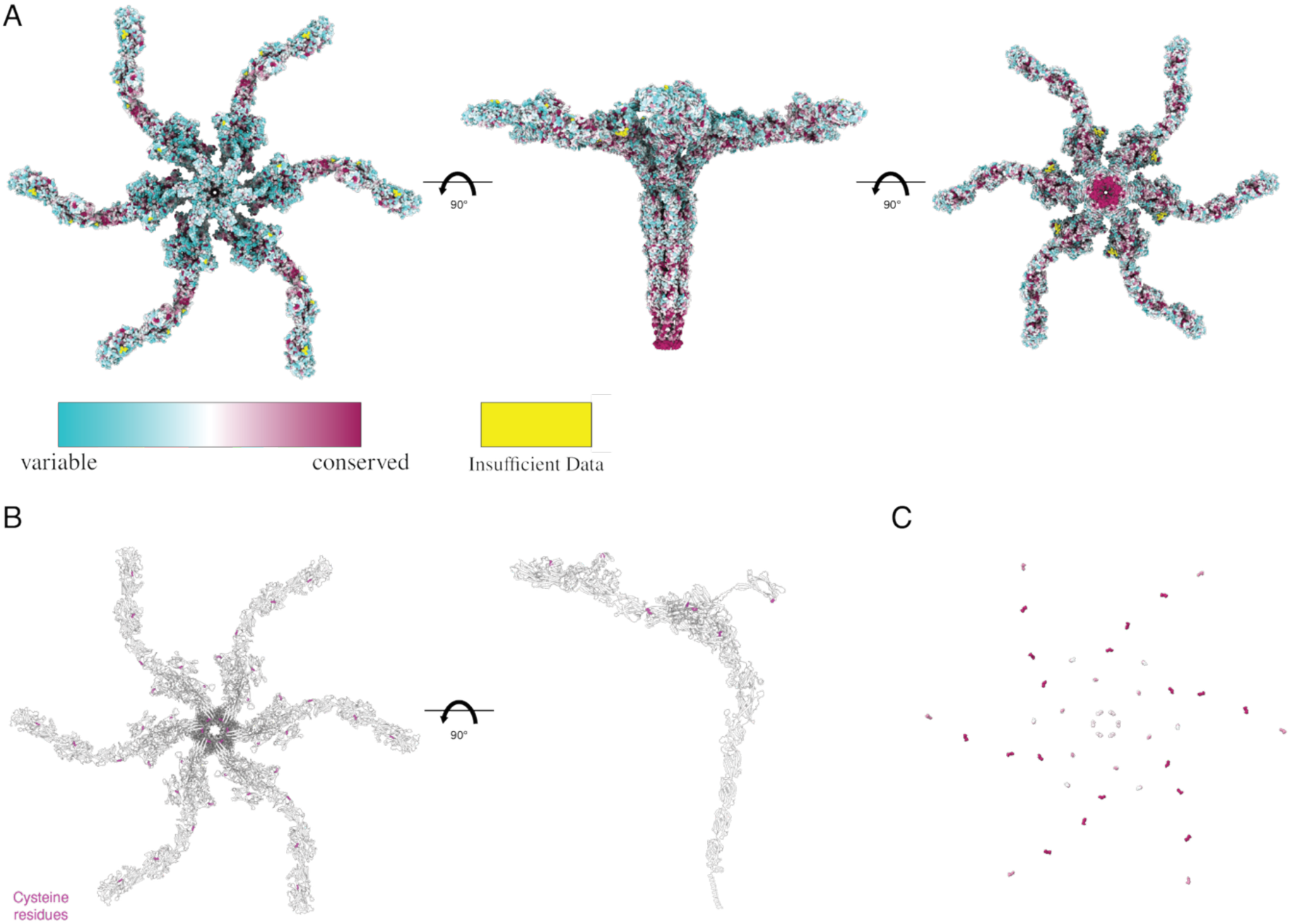
– Sequence conservation of candidate SLPs within the family Thermoproteaceae. **A**) Top (left, viewed from outside the cell) side (middle, viewed from the membrane) and bottom (right, viewed from the cytosol) representation of the PySLP hexamer coloured according to the conservation score derived from multiple alignment of sequence encoded by *Pyrobaculum* and *Thermoproteus* species. **B**) Cysteine residues potentially forming disulfide bridges in the lattice domains of PySLP. The cysteines are shown as magenta spheres on the hexamer (left, top view) and on the monomer (right, side view). **C**) PySLP cysteines coloured according to their conservation score, shown as spheres from a top view.

**Supplementary Fig. 11.**
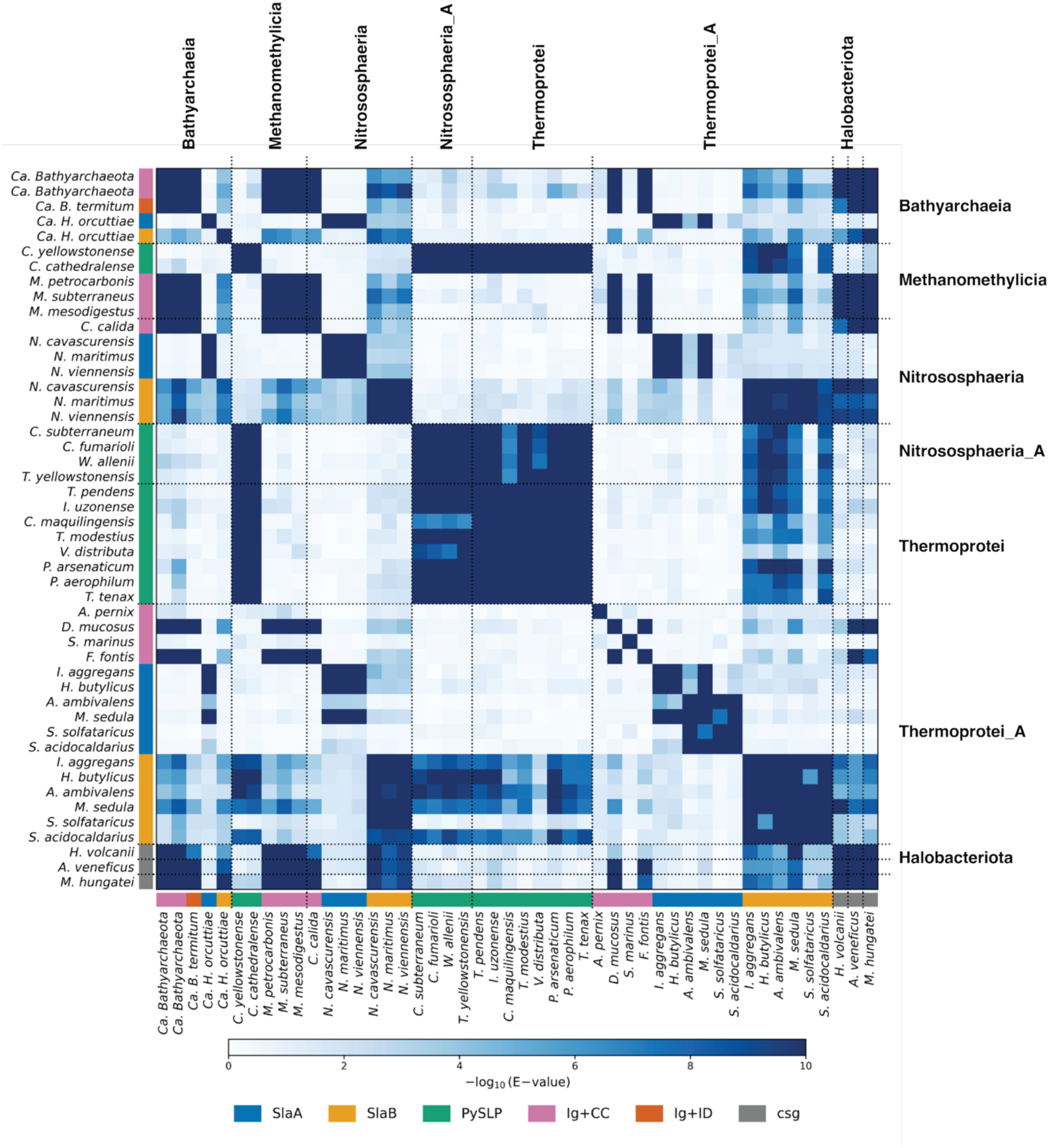
– All-against-all HMM-HMM similarity matrix of SLPs. Sequence similarity comparison of experimentally characterised and putative SLPs from different classes of Thermoproteota, together with reference SLPs from Halobacteriota. Pairwise similarities are shown as −log10(E-values), with darker colours indicating stronger similarity. Coloured bars indicate the different SLP architectural groups, and dotted lines delineate taxonomic groups. Ig, Ig-like domain; CC, coiled coil; ID, intrinsically disordered; csg, cell surface glycoprotein.

**Supplementary Fig. 12.**
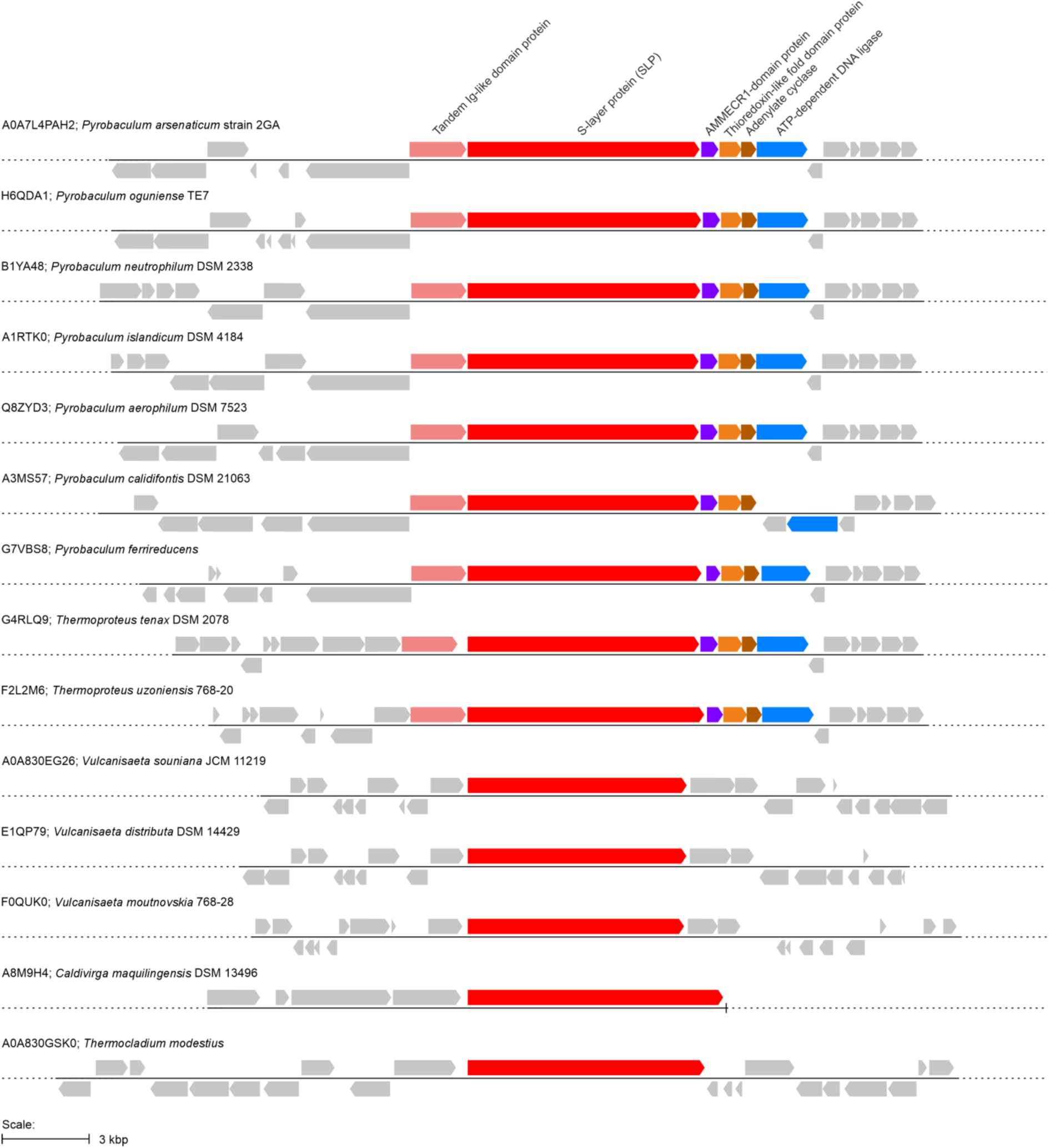
– Genomic neighbourhood of the SLP genes in the order *Thermoproteales*. Homologous genes in the putative conserved operon including the SLP gene are shown as arrows with matching colours. Other genes are shown in grey. Each locus is identified with the UniProt accession number of the corresponding SLP gene (red), followed by the name of the species.

**Supplementary Fig. 13.**
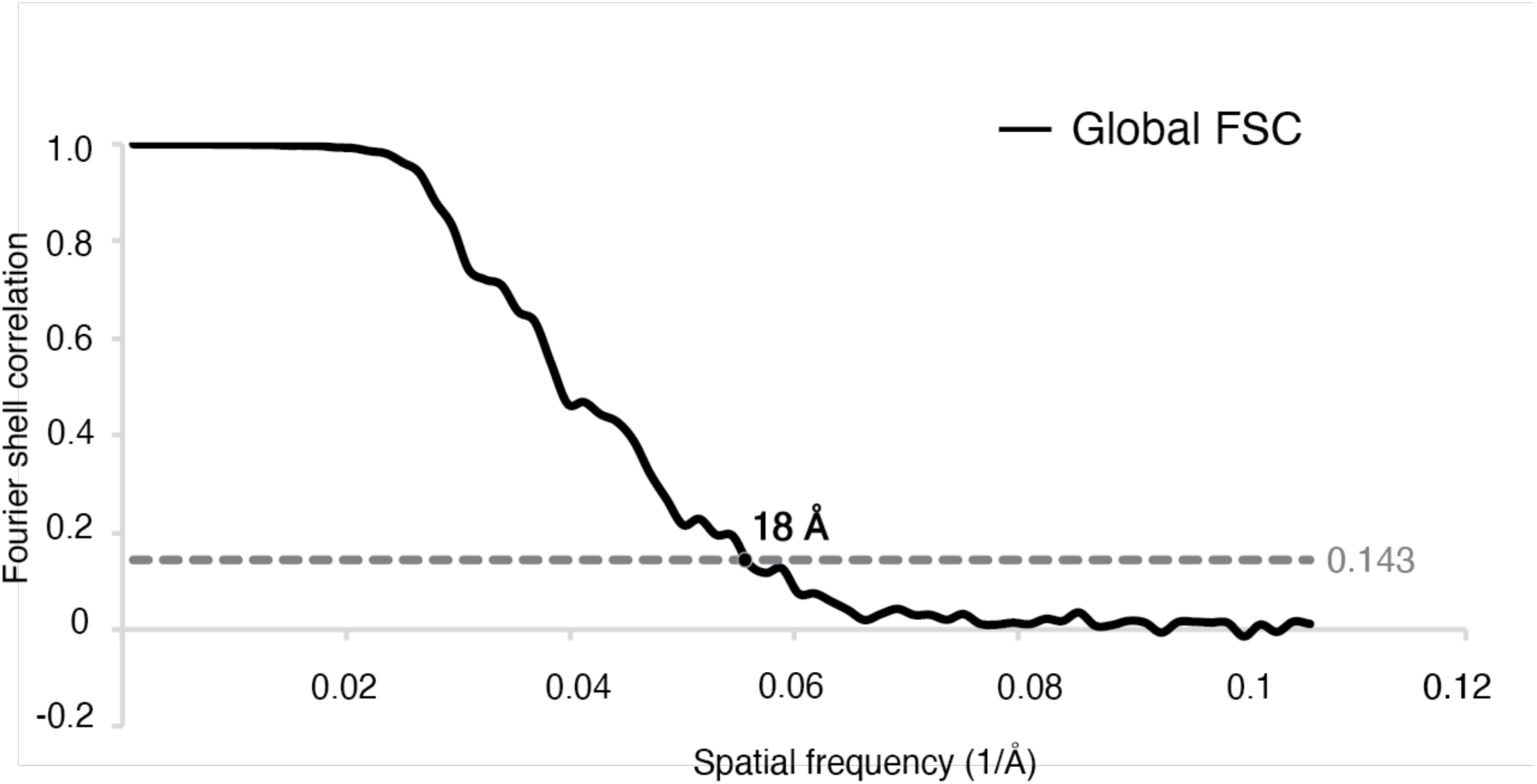
– Resolution estimation of the cryo-ET STA map. Global Fourier shell correlation (FSC) resolution estimation of the *in situ* STA map.

**Supplementary Movie 1 – Representative *P. arsenaticum* cellular tomogram.**

Description: The tomogram was CTF corrected and reconstructed using RELION5.0 and subsequently denoised using Cryo-CARE. Sequential Z-slices through the tomogram are shown. Scale bar – 200 nm.

**Supplementary Movie 2 – Representative *P. oguniense* cellular tomogram.**

Description: The tomogram was reconstructed using SIRT implemented in Tomo3D. Sequential Z-slices through the tomogram are shown. Scale bar – 200 nm.

**Supplementary Movie 3 – Molecular architecture of PySLP.**

Description: PySLP cryo-EM map shown in grey, zoom-in onto the PySLP monomer coloured rainbow, highlighting the lattice forming domain, crown, pillar and transmembrane helix. The cryo-EM map is shown again in transparency and with the fitted PySLP hexamer, with each monomer coloured separately, noting the pillar and crown formed at it centre. Zoom-out showing the fit of the surrounding six hexamers into the transparent cryo-EM map, constructing the PySLP lattice. Zoom-in onto the trimeric interface and N-terminal stabilising interface.

**Supplementary Table 1:**
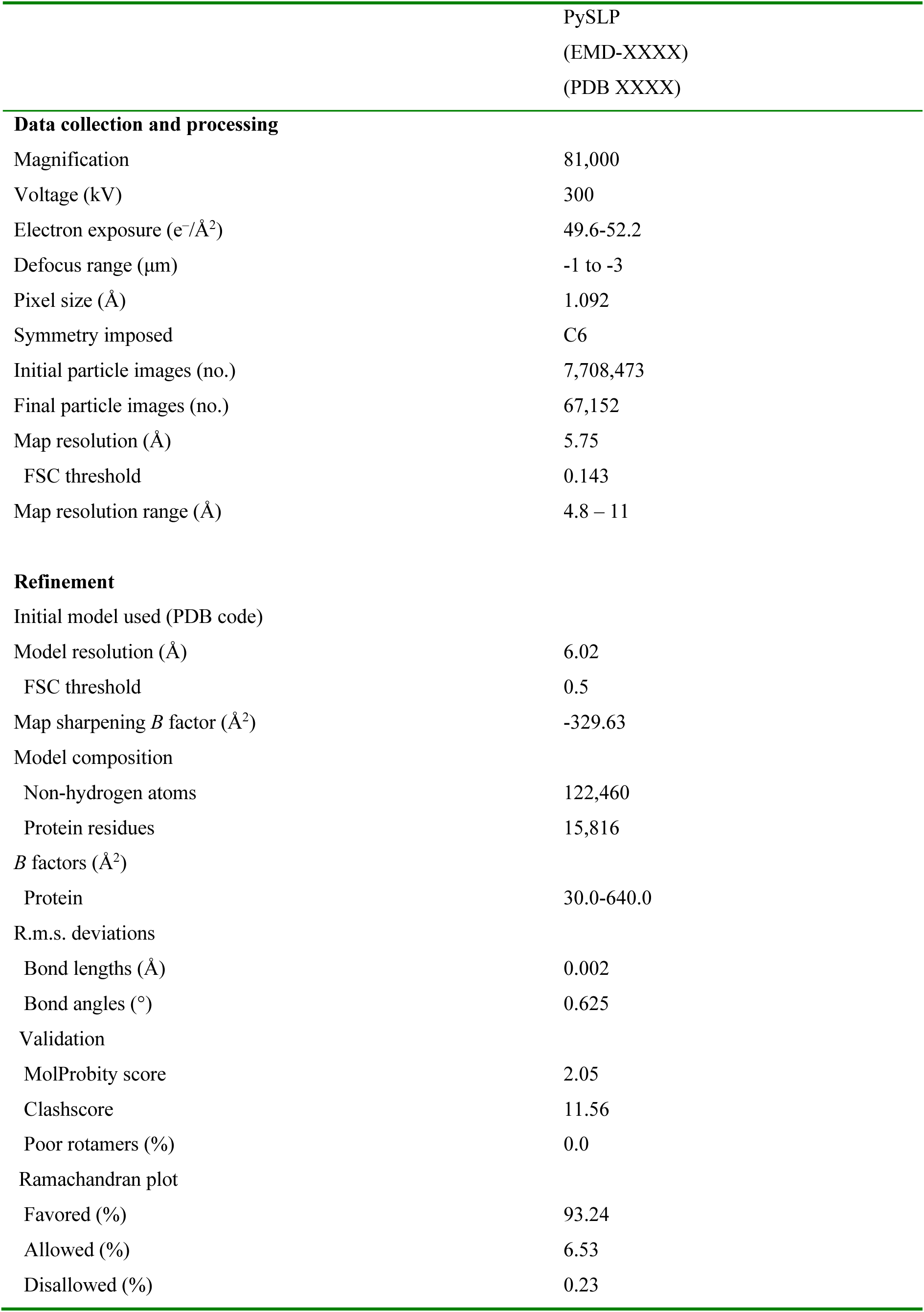
Cryo-EM data collection, refinement and validation statistics.

|  |  |
| --- | --- |
|  | PySLP<br>(EMD-XXXX)<br>(PDB XXXX) |
| **Data collection and processing** |  |
| Magnification | 81,000 |
| Voltage (kV) | 300 |
| Electron exposure (e<sup>-</sup>/Å<sup>2</sup>) | 49.6-52.2 |
| Defocus range (µm) | -1 to -3 |
| Pixel size (Å) | 1.092 |
| Symmetry imposed | C6 |
| Initial particle images (no.) | 7,708,473 |
| Final particle images (no.) | 67,152 |
| Map resolution (Å) | 5.75 |
| FSC threshold | 0.143 |
| Map resolution range (Å) | 4.8 – 11 |
| **Refinement** |  |
| Initial model used (PDB code) |  |
| Model resolution (Å) | 6.02 |
| FSC threshold | 0.5 |
| Map sharpening *B* factor (Å<sup>2</sup>) | -329.63 |
| Model composition |  |
| Non-hydrogen atoms | 122,460 |
| Protein residues | 15,816 |
| *B* factors (Å<sup>2</sup>) |  |
| Protein | 30.0-640.0 |
| R.m.s. deviations |  |
| Bond lengths (Å) | 0.002 |
| Bond angles (°) | 0.625 |
| Validation |  |
| MolProbity score | 2.05 |
| Clashscore | 11.56 |
| Poor rotamers (%) | 0.0 |
| Ramachandran plot |  |
| Favored (%) | 93.24 |
| Allowed (%) | 6.53 |
| Disallowed (%) | 0.23 |

**Supplementary Table 2:** Protein accessions from HHpred proteomic searches of Thermoproteota and selected Halobacteriota.

| Class | Organism | Protein accession (UniProt/UniRef/NCBI/GTDB) |
| --- | --- | --- |
| **Bathyarchaeia** | *Ca. Bathyarchaeota archaeon* | UPI000FF5155F; UPI000FF14A7B |
|  | *Ca. Bathyarchaeum termitum* | MDR0373548.1 |
|  | *Ca. Hecatella orcuttiae* | WP_309492968.1; WP_309492970.1 |
| **Methanomethylica** | *Ca. Culexarchaeum yellowstonense* | UPI0044657478 |
|  | *Ca. Culexmicrobium cathedralense* | MCS7385630.1 |
|  | *Ca. Methanosuratincola petrocarbonis* | A0A7J3UZ30 |
|  | *Ca. Methanosuratincola subterraneus* | A0A444L8N0 |
|  | *Methanomethylicus mesodigestus* | GTDB: MAGS01000030.1_118 |
| **Nitrososphaeria** | *Conexivisphaera calida* | A0A4P2VKM0 |
|  | *Ca. Nitrosocaldus cavascurensis* | A0A2K5AP30; A0A2K5AP17 |
|  | *Nitrosopumilus maritimus* | A9A4Y9; A9A4Y8 |
|  | *Nitrososphaera viennensis* | A0A060HS03; A0A060HR06 |
| **Nitrososphaeria_A** | *Ca. Caldarchaeum subterraneum* | BAJ47616.1 |
|  | *Ca. Calditenuis fumarioli* | MDJ0274653.1 |
|  | *Ca. Wolframiraptor allenii* | MCL7393743.1 |
|  | *Ca. Terraquivivens yellowstonensis* | MCL7394657.1 |
| **Thermoprotei** | *Thermofilum pendens* | A1RY64 |
|  | *Infirmifilum uzonense* | A0A0F7FI64 |
|  | *Caldivirga maquilingensis* | A8M9H4 |
|  | *Thermocladium modestius* | WP_229657656.1 |
|  | *Vulcanisaeta distributa* | E1QP79 |
|  | *Pyrobaculum arsenaticum* | A0A7L4PAH2 |
|  | *Pyrobaculum aerophilum* | Q8ZYD3 |
|  | *Thermoproteus tenax* | G4RLQ9 |
| **Thermoprotei_A** | *Aeropyrum pernix* | Q9YEG7 |
|  | *Desulfurococcus mucosus* | E8R795 |
|  | *Staphylothermus marinus* | Q54436 |
|  | *Fervidicoccus fontis* | I0A0F7 |
|  | *Ignisphaera aggregans* | E0SPF1; E0SPF2 |
|  | *Hyperthermus butylicus* | A2BLH8; A2BLH9 |
|  | *Acidianus ambivalens* | B1GT61; B1GT62 |
|  | *Metallosphaera sedula* | A4YHQ8; A4YHQ9 |
|  | *Saccharolobus solfataricus* | Q980C7; Q980C6 |
|  | *Sulfolobus acidocaldarius* | Q4J6E5; Q4J6E6 |
| **Halobacteriota** | *Archaeoglobus veneficus* | F2KQ80 |
|  | *Haloferax volcanii* | P25062 |
|  | *Methanospirillum hungatei* | Q2FTS3 |

## References

1. Sára, M. & Sleytr, U. B. S-Layer Proteins. J. Bacteriol. 182, 859 (2000).

2. Isbilir, B., von Kügelgen, A., Alva, V. & Bharat, T. A. M. Assembly, architecture and functional roles of microbial surface layers. Nat. Rev. Microbiol. 24, 344–358 (2025).

3. Fagan, R. P. & Fairweather, N. F. Biogenesis and functions of bacterial S-layers. Nat. Rev. Microbiol. 12, 211–222 (2014).

4. Bharat, T. A. M., von Kügelgen, A. & Alva, V. Molecular Logic of Prokaryotic Surface Layer Structures. Trends Microbiol. 29, 405 (2021).

5. Sleytr, U. B., Schuster, B., Egelseer, E. M. & Pum, D. S-layers: Principles and applications. FEMS Microbiol. Rev. 38, 823–864 (2014).

6. Albers, S. V. & Meyer, B. H. The archaeal cell envelope. Nat. Rev. Microbiol. 9, 414– 426 (2011).

7. Foo, S., Caspy, I., Cezanne, A., Bharat, T. A. M. & Baum, B. A self-assembling surface layer flattens the cytokinetic furrow to aid cell division in an archaeon. Proceedings of the National Academy of Sciences 122, e2501044122 (2025).

8. von Kügelgen, A. et al. Membraneless channels sieve cations in ammonia-oxidizing marine archaea. Nature 630, 230–236 (2024).

9. Mader, C., Küpcü, S., Sára, M. & Sleytr, U. B. Stabilizing effect of an S-layer on liposomes towards thermal or mechanical stress. Biochimica et Biophysica Acta (BBA) - Biomembranes 1418, 106–116 (1999).

10. Engelhardt, H. Mechanism of osmoprotection by archaeal S-layers: A theoretical study. J. Struct. Biol. 160, 190–199 (2007).

11. Schuster, B. & Sleytr, U. B. The effect of hydrostatic pressure on S-layer-supported lipid membranes. Biochimica et Biophysica Acta (BBA) - Biomembranes 1563, 29–34 (2002).

12. Rados, T. et al. Tissue-like multicellular development triggered by mechanical compression in archaea. Science (1979). 388, 109–115 (2025).

13. Tamir, A. & Eichler, J. N-Glycosylation is important for proper *Haloferax volcanii* S-layer stability and function. Appl. Environ. Microbiol. 83, (2017).

14. Grill-Walcher, S. & Schäffer, C. A new age in structural S-layer biology: Experimental and in silico milestones. Journal of Biological Chemistry 301, 110205 (2025).

15. van Wolferen, M. et al. Species-Specific Recognition of Sulfolobales Mediated by UV-Inducible Pili and S-Layer Glycosylation Patterns. mBio 11, e03014–19 (2020).

16. Vogel, C., Berzuini, C., Bashton, M., Gough, J. & Teichmann, S. A. Supra-domains: Evolutionary Units Larger than Single Protein Domains. J. Mol. Biol. 336, 809–823 (2004).

17. Bahl, H. et al. IV. Molecular biology of S-layers. FEMS Microbiol. Rev. 20, 47–98 (1997).

18. Hagen, K. E., Guan, L. L., Tannock, G. W., Korver, D. R. & Allison, G. E. Detection, Characterization, and In Vitro and In Vivo Expression of Genes Encoding S-Proteins in Lactobacillus gallinarum Strains Isolated from Chicken Crops. Appl. Environ. Microbiol. 71, 6633 (2005).

19. Rothschild, L. J. & Mancinelli, R. L. Life in extreme environments. Nature 409, 1092– 1101 (2001).

20. Auernik, K. S., Cooper, C. R. & Kelly, R. M. Life in hot acid: Pathway analyses in extremely thermoacidophilic archaea. Curr. Opin. Biotechnol. 19, 445 (2008).

21. Siliakus, M. F., van der Oost, J. & Kengen, S. W. M. Adaptations of archaeal and bacterial membranes to variations in temperature, pH and pressure. Extremophiles 21, 651–670 (2017).

22. Vieille, C. & Zeikus, G. J. Hyperthermophilic Enzymes: Sources, Uses, and Molecular Mechanisms for Thermostability. Microbiology and Molecular Biology Reviews 65, 1 (2001).

23. Jorda, J. & Yeates, T. O. Widespread Disulfide Bonding in Proteins from Thermophilic Archaea. Archaea 2011, 409156 (2011).

24. Wildhaber, I. & Baumeister, W. The cell envelope of *Thermoproteus tenax*: three-dimensional structure of the surface layer and its role in shape maintenance. EMBO J. 6, 1475 (1987).

25. Messner, P., Pum, D., Sara, M., Stetter, K. O. & Sleytr, U. B. Ultrastructure of the cell envelope of the archaebacteria *Thermoproteus tenax* and *Thermoproteus neutrophilus*. J. Bacteriol. 166, 1046 (1986).

26. Phipps, B. M., Huber, R. & Baumeister, W. The cell envelope of the hyperthermophilic archaebacterium *Pyrobaculum organotrophum* consists of two regularly arrayed protein layers: three-dimensional structure of the outer layer. Mol. Microbiol. 5, 253–265 (1991).

27. Phipps, B. M., Engelhardt, H., Huber, R. & Baumeister, W. Three-dimensional structure of the crystalline protein envelope layer of the hyperthermophilic archaebacterium *Pyrobaculum islandicum*. J. Struct. Biol. 103, 152–163 (1990).

28. Dürr, R., Hegerl, R., Volker, S., Santarius, U. & Baumeister, W. Three-dimensional reconstruction of the surface protein of *Pyrodictium brockii*: Comparing two image processing strategies. J. Struct. Biol. 106, 181–190 (1991).

29. Gambelli, L. et al. Structure of the two-component S-layer of the archaeon *Sulfolobus acidocaldarius*. Elife 13, (2024).

30. Huber, R., Sacher, M., Vollmann, A., Huber, H. & Rose, D. Respiration of arsenate and selenate by hyperthermophilic archaea. Syst. Appl. Microbiol. 23, 305–314 (2000).

31. Jung, J. H. et al. Broad substrate specificity of a hyperthermophilic α-glucosidase from *Pyrobaculum arsenaticum*. Food Sci. Biotechnol. 25, 1665–1669 (2016).

32. Rensen, E., Krupovic, M. & Prangishvili, D. Mysterious hexagonal pyramids on the surface of Pyrobaculum cells. Biochimie 118, 365–367 (2015).

33. Rensen, E. et al. A virus of hyperthermophilic archaea with a unique architecture among DNA viruses. Proc. Natl. Acad. Sci. U. S. A. 113, 2478–2483 (2016).

34. Jay, Z. J., et al. *Pyrobaculum yellowstonensis* Strain WP30 Respires on Elemental Sulfur and/or Arsenate in Circumneutral Sulfidic Geothermal Sediments of Yellowstone National Park. Appl. Environ. Microbiol. 81, 5907 (2015).

35. Slobodkina, G. B., Lebedinsky, A. V., Chernyh, N. A., Bonch-Osmolovskaya, E. A. & Slobodkin, A. I. *Pyrobaculum ferrireducens* sp. nov., a hyperthermophilic Fe(III)-, selenate- and arsenate reducing crenarchaeon isolated from a hot spring. Int. J. Syst. Evol. Microbiol. 65, 851–856 (2015).

36. Seeholzer, T. et al. A Next-Generation qPlus-Sensor-Based AFM Setup: Resolving Archaeal S-Layer Protein Structures in Air and Liquid. Journal of Physical Chemistry B 127, 6949–6957 (2023).

37. Jumper, J. et al. Highly accurate protein structure prediction with AlphaFold. Nature 596, 583–589 (2021).

38. Mirdita, M. et al. ColabFold: making protein folding accessible to all. Nat. Methods 19, 679–682 (2022).

39. von Kügelgen, A., Alva, V. & Bharat, T. A. M. Complete atomic structure of a native archaeal cell surface. Cell Rep. 37, (2021).

40. von Kügelgen, A. et al. In Situ Structure of an Intact Lipopolysaccharide-Bound Bacterial Surface Layer. Cell 180, 348 (2020).

41. Isbilir, B., Yeates, A., Alva, V. & Bharat, T. A. M. Mapping the ultrastructural topology of the corynebacterial cell surface. PLoS Biol. 23, e3003130 (2025).

42. Sogues, A. et al. Cryo-EM structure and polar assembly of the PS2 S-layer of *Corynebacterium glutamicum*. Proc. Natl. Acad. Sci. U. S. A. 122, e2426928122 (2025).

43. Teufel, F. et al. SignalP 6.0 predicts all five types of signal peptides using protein language models. Nat. Biotechnol. 40, 1023–1025 (2022).

44. Abramson, J. et al. Accurate structure prediction of biomolecular interactions with AlphaFold 3. Nature 630, 493–500 (2024).

45. von Kügelgen, A. et al. Interdigitated immunoglobulin arrays form the hyperstable surface layer of the extremophilic bacterium *Deinococcus radiodurans*. Proc. Natl. Acad. Sci. U. S. A. 120, e2215808120 (2023).

46. Hallgren, J. et al. DeepTMHMM predicts alpha and beta transmembrane proteins using deep neural networks. bioRxiv 10.1101/2022.04.08.487609 (2022) doi:10.1101/2022.04.08.487609.

47. Volkl, P., et al. *Pyrobaculum aerophilum* sp. nov., a novel nitrate-reducing hyperthermophilic archaeum. Appl. Environ. Microbiol. 59, 2918 (1993).

48. Gambelli, L. et al. Architecture and modular assembly of Sulfolobus S-layers revealed by electron cryotomography. Proc. Natl. Acad. Sci. U. S. A. 116, 25278–25286 (2019).

49. Wang, F., Cvirkaite-Krupovic, V., Krupovic, M. & Egelman, E. H. Archaeal bundling pili of *Pyrobaculum calidifontis* reveal similarities between archaeal and bacterial biofilms. Proc. Natl. Acad. Sci. U. S. A. 119, e2207037119 (2022).

50. Wang, F. et al. The structures of two archaeal type IV pili illuminate evolutionary relationships. Nat. Commun. 11, 1–10 (2020).

51. Bharat, T. A. M. et al. Structure of the hexagonal surface layer on *Caulobacter crescentus* cells. Nat. Microbiol. 2, 17059-(2017).

52. Baranova, E. et al. SbsB structure and lattice reconstruction unveil Ca2+ triggered S-layer assembly. Nature 487, 119–122 (2012).

53. Holm, L., Laiho, A., Törönen, P. & Salgado, M. DALI shines a light on remote homologs: One hundred discoveries. Protein Science 32, e4519 (2023).

54. van Kempen, M. et al. Fast and accurate protein structure search with Foldseek. Nat. Biotechnol. 42, 243–246 (2023).

55. Zimmermann, L. et al. A Completely Reimplemented MPI Bioinformatics Toolkit with a New HHpred Server at its Core. J. Mol. Biol. 430, 2237–2243 (2018).

56. Wang, H. et al. Composition and in situ structure of the *Methanospirillum hungatei* cell envelope and surface layer. Sci. Adv. 10, 8596 (2024).

57. Lanzoni-Mangutchi, P. et al. Structure and assembly of the S-layer in *C. difficile*. Nat. Commun. 13, 970 (2022).

58. Björklund, Å. K., Ekman, D. & Elofsson, A. Expansion of Protein Domain Repeats. PLoS Comput. Biol. 2, e114 (2006).

59. Altschul, S. F., Gish, W., Miller, W., Myers, E. W. & Lipman, D. J. Basic local alignment search tool. J. Mol. Biol. 215, 403–410 (1990).

60. Itoh, T. The family thermoproteaceae. in The Prokaryotes: Other Major Lineages of Bacteria and The Archaea (eds. Rosenberg, E., DeLong, E. F., Lory, S., Stackebrandt, E. & Thompson, F.) 389–401 (Springer-Verlag Berlin Heidelberg, 2014). doi:10.1007/978-3-642-38954-2_330.

61. Siebers, B. et al. The Complete Genome Sequence of Thermoproteus tenax: A Physiologically Versatile Member of the Crenarchaeota. PLoS One 6, e24222 (2011).

62. Peters, J. et al. Tetrabrachion: A Filamentous Archaebacterial Surface Protein Assembly of Unusual Structure and Extreme Stability. J. Mol. Biol. 245, 385–401 (1995).

63. Johnston, E., Isbilir, B., Alva, V., Bharat, T. A. M. & Doye, J. P. K. Punctuated and continuous structural diversity of S-layers across the prokaryotic tree of life. bioRxiv 10.1101/2024.05.28.596244 (2024) doi:10.1101/2024.05.28.596244.

64. Sogues, A. et al. Architecture of the Sap S-layer of *Bacillus anthracis* revealed by integrative structural biology. Proc. Natl. Acad. Sci. U. S. A. 121, e2415351121 (2024).

65. Sivabalasarma, S., van Wolferen, M. & Albers, S. V. Biogenesis, function and evolution of the archaeal S-layer. Curr. Opin. Cell Biol. 95, 102534 (2025).

66. Beeby, M. et al. The Genomics of Disulfide Bonding and Protein Stabilization in Thermophiles. PLoS Biol. 3, e309 (2005).

67. Krupovic, M., White, M. F., Forterre, P. & Prangishvili, D. Postcards from the Edge: Structural Genomics of Archaeal Viruses. in Advances in Virus Research vol. 82 33–62 (Academic Press, 2012).

68. Engelhardt, H. Are S-layers exoskeletons? The basic function of protein surface layers revisited. J. Struct. Biol. 160, 115–124 (2007).

69. Baquero, D. P. et al. New virus isolates from Italian hydrothermal environments underscore the biogeographic pattern in archaeal virus communities. ISME J. 14, 1821 (2020).

70. Sako, Y., Nunoura, T. & Uchida, A. *Pyrobaculum oguniense* sp. nov., a novel facultatively aerobic and hyperthermophilic archaeon growing at up to 97 °C. Int. J. Syst. Evol. Microbiol. 51, 303–309 (2001).

71. Mastronarde, D. N. Automated electron microscope tomography using robust prediction of specimen movements. J. Struct. Biol. 152, 36–51 (2005).

72. Zivanov, J. et al. A Bayesian approach to single-particle electron cryo-tomography in RELION-4.0. Elife 11, (2022).

73. Zheng, S. Q. et al. MotionCor2: anisotropic correction of beam-induced motion for improved cryo-electron microscopy. Nat. Methods 14, 331–332 (2017).

74. Rohou, A. & Grigorieff, N. CTFFIND4: Fast and accurate defocus estimation from electron micrographs. J. Struct. Biol. 192, 216–221 (2015).

75. Bepler, T. et al. Positive-unlabeled convolutional neural networks for particle picking in cryo-electron micrographs. Nat. Methods 16, 1153–1160 (2019).

76. Punjani, A., Rubinstein, J. L., Fleet, D. J. & Brubaker, M. A. cryoSPARC: algorithms for rapid unsupervised cryo-EM structure determination. Nat. Methods 14, 290–296 (2017).

77. Punjani, A., Zhang, H. & Fleet, D. J. Non-uniform refinement: adaptive regularization improves single-particle cryo-EM reconstruction. Nat. Methods 17, 1214–1221 (2020).

78. Zivanov, J., Nakane, T. & Scheres, S. H. W. Estimation of high-order aberrations and anisotropic magnification from cryo-EM data sets in RELION-3.1. IUCrJ 7, 253–267 (2020).

79. Baquero, D. P. et al. Biogenesis of DNA-carrying extracellular vesicles by the dominant human gut methanogenic archaeon. Nat. Commun. 16, 5093 (2025).

80. Cox, J. et al. Andromeda: A Peptide Search Engine Integrated into the MaxQuant Environment. J. Proteome Res. 10, 1794–1805 (2011).

81. Tyanova, S., Temu, T. & Cox, J. The MaxQuant computational platform for mass spectrometry-based shotgun proteomics. Nat. Protoc. 11, 2301–2319 (2016).

82. Emsley, P., Lohkamp, B., Scott, W. G. & Cowtan, K. Features and development of Coot. Acta Crystallogr. D Biol. Crystallogr. 66, 486–501 (2010).

83. Yamashita, K., Wojdyr, M., Long, F., Nicholls, R. A. & Murshudov, G. N. GEMMI and Servalcat restrain REFMAC5. Acta Crystallogr. D Struct. Biol. 79, 368–373 (2023).

84. Liebschner, D. et al. Macromolecular structure determination using X-rays, neutrons and electrons: Recent developments in Phenix. Acta Crystallogr. D Struct. Biol. 75, 861– 877 (2019).

85. Goddard, T. D. et al. UCSF ChimeraX: Meeting modern challenges in visualization and analysis. Protein Science 27, 14–25 (2018).

86. Kremer, J. R., Mastronarde, D. N. & McIntosh, J. R. Computer visualization of three-dimensional image data using IMOD. J. Struct. Biol. 116, 71–76 (1996).

87. Agulleiro, J. I. & Fernandez, J. J. Tomo3D 2.0--exploitation of advanced vector extensions (AVX) for 3D reconstruction. J. Struct. Biol. 189, 147–152 (2015).

88. Fernandez, J. J., Li, S., Bharat, T. A. M. & Agard, D. A. Cryo-tomography tilt-series alignment with consideration of the beam-induced sample motion. J. Struct. Biol. 202, 200–209 (2018).

89. Burt, A. et al. An image processing pipeline for electron cryo-tomography in RELION-5. FEBS Open Bio 14, 1788–1804 (2024).

90. Buchholz, T. O., Jordan, M., Pigino, G. & Jug, F. Cryo-CARE: Content-Aware Image Restoration for Cryo-Transmission Electron Microscopy Data. Proceedings - International Symposium on Biomedical Imaging 2019-April, 502–506 (2018).

91. Chaillet, M. L., Roet, S., Veltkamp, R. C. & Förster, F. pytom-match-pick: A tophat-transform constraint for automated classification in template matching. J. Struct. Biol. X 11, 100125 (2025).

92. Ermel, U. H., Arghittu, S. M. & Frangakis, A. S. ArtiaX: An electron tomography toolbox for the interactive handling of sub-tomograms in UCSF ChimeraX. Protein Science 31, e4472 (2022).

93. Goldfarb, T. et al. NCBI RefSeq: reference sequence standards through 25 years of curation and annotation. Nucleic Acids Res. 53, D243 (2024).

94. Sayers, E. W. et al. Database resources of the National Center for Biotechnology Information in 2026. Nucleic Acids Res. 54, D20 (2025).

95. Steinegger, M. et al. HH-suite3 for fast remote homology detection and deep protein annotation. BMC Bioinformatics 20, 473 (2019).

96. Mirdita, M. et al. Uniclust databases of clustered and deeply annotated protein sequences and alignments. Nucleic Acids Res. 45, D170 (2016).

97. Yamashita, K., Palmer, C. M., Burnley, T. & Murshudov, G. N. Cryo-EM single-particle structure refinement and map calculation using Servalcat. Acta Crystallogr. D Struct. Biol. 77, 1282–1291 (2021).

